# Interstitium-mimicking porous alveolar membranes enable physiologic aerosol transport and distinct acute-chronic lung injury responses

**DOI:** 10.64898/2026.06.01.729429

**Authors:** Jae-Won Choi, Sheng Zhang, Abbas Jalili, August Kohls, Woo-Youl Maeng, Sikandar Azam, Wu Liu, Barbie Varghese, Yixin Zhao, Xi Ren, Shimin Liu, Si-Yang Zheng

## Abstract

Barrier membranes govern transport and mechanochemical coupling in lung-on-chip systems but typically exhibit low open porosity, limited pore interconnectivity, and diffusion distances exceeding native thin septal regions. An interstitium-mimicking, alveolus-shaped poly(ε-caprolactone) membrane is developed using dual-templated nonsolvent-induced phase separation followed by controlled enzymatic pore enlargement. The resulting architecture achieves ∼40% total porosity with 97% pore interconnectivity and incorporates a locally thinned dome region (∼2.5 µm). This structure sustains cyclic deformation while increasing oxygen diffusivity fivefold compared with conventional Transwell® membranes under both acellular and epithelial-endothelial co-culture conditions. Integrated into an air-liquid interface platform, the membrane enables direct aerosol deposition and quantitative interrogation of cross-barrier mass transfer. Using carbonaceous nanoscale particulate matter as a model inhaled aerosol, controlled exposure induces dose-dependent oxidative, inflammatory, and genotoxic responses. Matched cumulative dose studies reveal distinct biological trajectories: acute high-dose exposure produces rapid cytotoxic stress and barrier disruption, whereas chronic low-dose exposure preserves viability yet promotes sustained DNA repair and genome-maintenance programs. Compartment-resolved analysis and therapeutic intervention further demonstrate the platform’s utility for spatial and translational interrogation of lung injury. By restoring physiologically relevant diffusion distance, interconnectivity, and strain responsiveness, the interstitium-mimicking membrane advances lung-on-chip design toward functional replication of alveolar transport dynamics for studying lung injury and barrier dysfunction.

## 1. Introduction

The alveolar air-blood barrier is a central functional unit in lung-on-chip systems, where epithelial-endothelial co-culture and cyclic deformation are used to emulate gas exchange and injury responses [1, 2]. Despite advances in microphysiological platforms, the barrier membrane frequently becomes the dominant rate-limiting element in these systems [3]. Commonly used membranes exhibit limited open pathways, incomplete pore interconnectivity, and thicknesses exceeding native thin septal regions, thereby imposing non-physiologic diffusion resistance and attenuating strain transmission across the interface. As a result, cross-barrier molecular exchange, epithelial-endothelial crosstalk, and aerosol transport dynamics are constrained by the artificial membrane rather than governed by intrinsic tissue behavior. Overcoming this membrane-imposed transport bottleneck is essential for faithfully replicating alveolar function *in vitro*.

In the native lung, alveolar septa achieve exceptional gas-exchange efficiency through ultrathin diffusion distances embedded within a compliant three-dimensional architecture. In thin septal regions of the native alveolus, epithelial and endothelial basement membranes are closely apposed, resulting in an air-blood barrier thickness of approximately 0.2-2 μm and thereby minimizing diffusion distance while maintaining structural integrity [4–6]. The intervening interstitial matrix forms a continuous and permeable scaffold that facilitates rapid exchange of gases, soluble mediators, and deposited particles between air-exposed epithelium and perfused endothelium [7, 8]. These transport processes occur under cyclic mechanical deformation during respiration, where repetitive strain dynamically alters local geometry and intercellular junctional tension. Together, high effective porosity, short diffusion distance, and mechanically stable deformation enable efficient transport while preserving barrier function, making them defining characteristics of the alveolar interface [9, 10]. Disruption of this finely tuned structure–transport–mechanics coupling contributes to a broad spectrum of pulmonary disorders, including pneumonia, chronic obstructive pulmonary disease, acute respiratory distress syndrome, pulmonary fibrosis, asthma, ventilation-induced injury, smoking-related disease, particulate matter exposure, and lung cancer, several of which remain leading causes of death worldwide [11].

*In vitro* models of the alveolar barrier provide important experimental control but incompletely reproduce this structure-transport coupling. Two-dimensional cultures lack three-dimensional curvature and strain-dependent mechanics [12], while Transwell® systems permit air-liquid interface (ALI) culture yet rely on flat, relatively impermeable membranes that impose non-physiologic diffusion resistance [13]. Organoids capture aspects of epithelial self-organization but form enclosed architectures that limit controlled aerosol delivery and perfusable endothelial interfaces [14, 15]. Lung-on-chip platforms introduce co-culture and cyclic deformation [16, 17]; however, many continue to employ membranes with low open area, limited pore interconnectivity, or thicknesses exceeding native thin septal regions, thereby constraining molecular exchange and attenuating epithelial-endothelial signaling under strain [1, 18–23]. Across these systems, the barrier membrane frequently becomes the dominant transport bottleneck, preventing faithful replication of alveolar permeability and particle translocation dynamics (Table S1) [19, 24–27]. Many lung-on-chip platforms use porous polydimethylsiloxane (PDMS) membranes ∼50 µm thick or porous polyethylene terephthalate (PET) or polycarbonate membranes ∼10 µm thick, both of which still differ from the native thin septal architecture [18–20, 26, 28, 29]. More recently, several in vitro alveolar models have incorporated electrospun ultrathin membrane mimics with thicknesses of ∼1-10 µm to better approximate the native air–blood barrier [30–33]. However, reducing membrane thickness alone does not fully recapitulate alveolar transport dynamics, as these systems generally lack the combination of high pore interconnectivity and three-dimensional alveolar geometry that characterizes the native septum.

To overcome membrane-imposed transport constraints, an interstitium-mimicking, alveolus-shaped porous membrane is developed as the structural core of an engineered alveolar interface. Fabricated from poly(ε-caprolactone) using dual-templated nonsolvent-induced phase separation followed by controlled enzymatic pore enlargement, the membrane achieves ∼40% total porosity with 97% pore interconnectivity and incorporates a locally thinned dome region (∼2.5 µm) that approaches thin septal dimensions while remaining mechanically stable under cyclic deformation. This architecture increases effective open pathways and reduces diffusion distance, resulting in approximately five-fold higher oxygen diffusivity than conventional Transwell® membranes under both acellular and epithelial–endothelial co-culture conditions. Integrated into an ALI epithelial-endothelial platform with controlled strain, the membrane enables direct aerosol delivery and quantitative interrogation of cross-barrier particle transport. The alveoli-on-chip platform is therefore used as a demonstration system to establish the functional consequences of the membrane architecture under physiologically relevant conditions.

Carbonaceous particulate matter, characteristic of combustion-derived and certain mining-associated aerosols, contribute substantially to global respiratory morbidity [34–38]. In occupational mining settings, prior investigations have often focused on micron-scale and silica-containing dust fractions using monolayer cultures and animal models [39–41]. By contrast, nanoscale organic-rich particles, which exhibit distinct physicochemical properties and preferential distal deposition [42], have been less frequently examined in systems that preserve an intact epithelial-endothelial barrier. Consequently, how such particles traverse the alveolar interface and how exposure pattern and mechanical strain influence downstream injury responses remain unclear. Using such nanoscale particulates as a model inhaled aerosol, quantifiable particle translocation across an intact epithelial-endothelial barrier is demonstrated, and matched cumulative dose studies reveal divergent biological trajectories: acute high-dose exposure induces rapid oxidative cytotoxicity and barrier disruption, whereas chronic low-dose exposure preserves viability yet promotes sustained DNA damage responses and genome-maintenance programs. Pharmacologic modulation further shows mitigation of key injury responses using antioxidant and anti-inflammatory agents. Together, these findings establish a structurally defined, transport-competent alveolar membrane that enables mechanistic investigation of inhalation-driven lung injury and barrier dysfunction.

## 2. Results

### 2.1 Interstitium-mimicking porous alveolar-shaped membrane architecture

The interstitium-mimicking membrane was engineered to recapitulate key structural and mechanical features of the native alveolar septum while minimizing membrane-imposed transport resistance. Design considerations were guided by three physiologically motivated criteria: (i) high vertical pore interconnectivity to facilitate cross-interface exchange, (ii) locally reduced thickness within dome regions to approximate thin septal diffusion distances, and (iii) sufficient mechanical compliance to sustain cyclic deformation within physiologic strain ranges without structural failure. To achieve this architecture, an alveolus-shaped porous membrane was fabricated from poly(ε-caprolactone) (PCL), selected for its mechanical compliance and biocompatibility under long-term epithelial–endothelial culture [43]. A dual-templated nonsolvent-induced phase separation strategy combined with controlled enzymatic pore enlargement was employed (Figure S1,S2). During solvent exchange, PCL precipitation and dendritic camphor crystallization jointly defined a vertically continuous pore network across the membrane thickness [44].

Pore architecture was modulated using two orthogonal parameters: camphor loading during casting and post-fabrication lipase treatment. Increasing camphor content from 0 to 250 wt% progressively increased surface porosity from 1.45% to 39.74%, accompanied by a reduction in Young’s modulus from 23.09 MPa to 6.14 MPa (Figure S3; Table S2), illustrating the expected trade-off between permeability and elastic stiffness. At the highest porogen ratios, membranes became mechanically fragile, therefore 150 wt% camphor was selected to preserve structural continuity while achieving high open area. Subsequent lipase treatment enabled controlled pore enlargement through partial ester cleavage of PCL [45, 46]. Under optimized lipase treatment conditions (0.05 mg mLD¹, 24 h), SEM analysis showed that the apical surface of the porous PCL membrane exhibited 23.4% porosity with an average pore diameter of ∼2.8 µm, whereas the basolateral surface reached 32.4% porosity with ∼5.4 µm pores (Figure 1A-C). Gravimetric analysis indicated that overall membrane porosity increased from 30.9% prior to lipase treatment to 36.8% after treatment (Figure 1D). Nano-computed tomography further confirmed 40.5% bulk porosity with 39.3% interconnected porosity (97% interconnectivity) (Figure 1E,F; Figure S4). Scanning electron microscopy confirmed vertically aligned pores with minimal dead-end structures (Figure 1G), establishing a highly connected transport pathway and closely resembling the native alveolar-capillary interface [47–51]. A locally thinned dome region of 2.64 µm connected by inter-alveolar regions (7.67 µm), thereby combining minimized diffusion distance with mechanical reinforcement (Figure 1H) [5]. To reproduce alveolar topology, the membrane was patterned into a 22 × 22 array of hemispherical units with a characteristic diameter of ∼200 µm, comparable to native alveolar dimensions (Figure 1I) [52]. Geometric analysis indicated that the flat inter-alveolar region accounted for approximately 87.5% of the total membrane surface area, whereas the curved alveolar dome region accounted for 12.5%, corresponding to a 2D:3D area ratio of approximately 7:1. This dome-shaped architecture introduces curvature and spatially heterogeneous strain fields during cyclic deformation while preserving thicker inter-alveolar support regions for structural stability.

**Figure 1.**
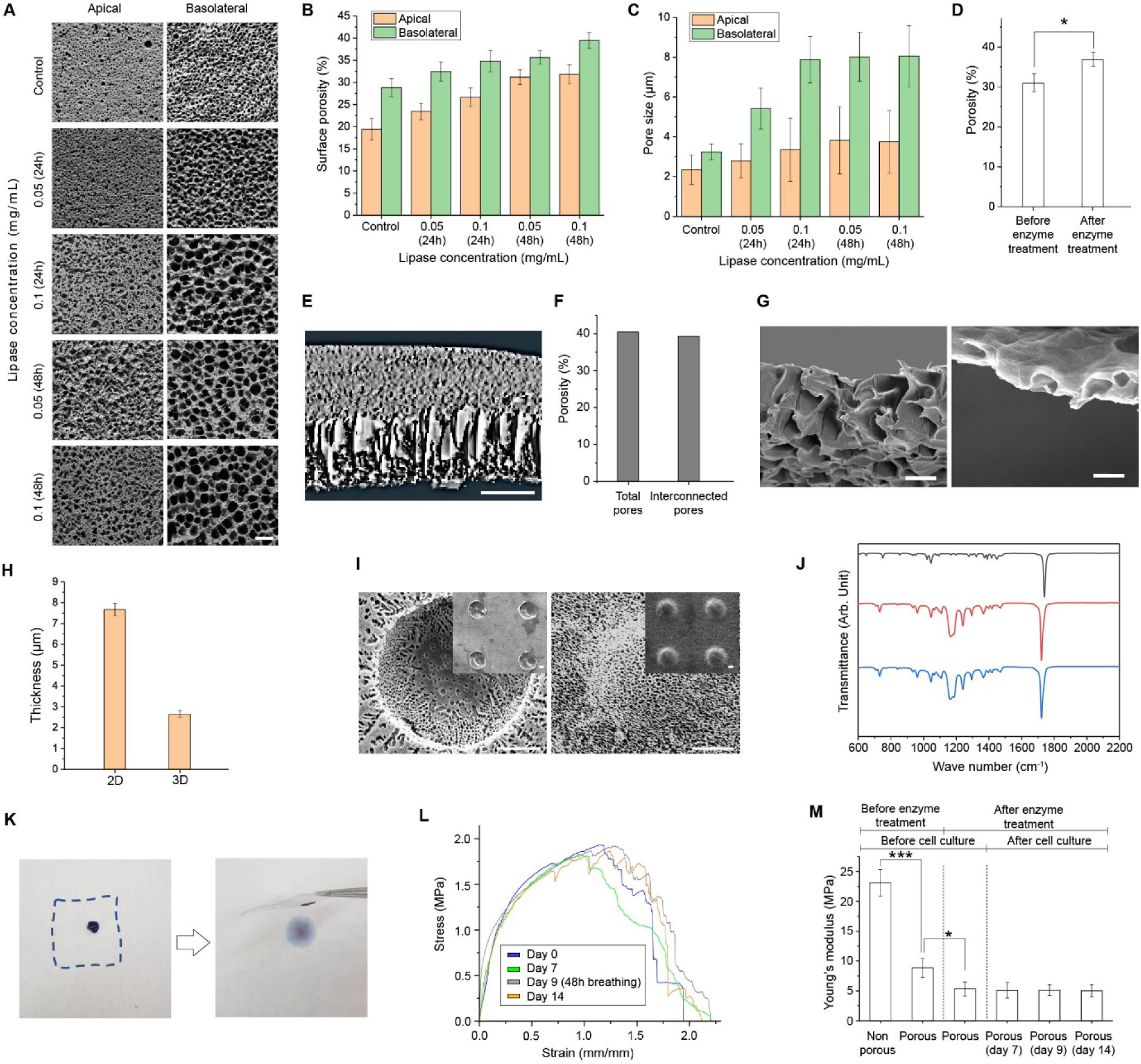
Structural and mechanical characterization of the interstitium-mimicking membrane. (**A**) Representative SEM images showing the surface morphology of porous PCL membranes on the apical and basolateral sides following lipase treatment at varying enzyme concentrations and exposure times. Scale bar: 10Dµm. (**B**) Quantified pore size on apical and basolateral surfaces as a function of lipase concentration and exposure time. (**C**) Quantitative analysis of apical and basolateral membrane surface porosity under varying enzyme concentrations and treatment times based on SEM image assessment. (**D**) Overall membrane porosity measured prior to and after enzymatic treatment by gravimetric analysis (n = 8). (**E**) Cross-sectional nano-CT image of the interstitium-mimicking membrane. Scale bar: 5 µm. (**F**) Total and interconnected porosity quantified from nano-CT analysis. (**G**) Cross-sectional SEM images of the membrane after lipase treatment, showing vertically interconnected pore structures in flat regions (left) and alveolus-shaped dome regions (right). Scale bar: 3Dµm. (**H**) Thickness comparison of the PCL membrane between flat regions (2D) and alveolus-shaped dome regions (3D), measured from cross-sectional SEM images (n = 4). (**I**) Top and bottom views of the alveolus-shaped membrane array with higher-magnification insets. Scale bar: 50 μm. (**J**) FT-IR spectra of camphor (black), interstitium-mimicking membrane (red), and pure PCL (blue), indicating successful removal of the porogen. (**K**) Optical images demonstrating through-thickness permeability by rapid wicking of aqueous dye across the membrane (dashed lines indicate membrane boundary). (**L**) Representative stress-strain curves of nonporous PCL, porous PCL, and lipase-treated interstitium-mimicking membranes at Days 0, 7, 9, and 14 of culture. (**M**) Mechanical characterization of nonporous PCL, porous PCL, and lipase-treated interstitium-mimicking membranes over 14 days of ALI culture (n = 5).

Fourier transform infrared spectroscopy confirmed successful removal of the camphor porogen following freeze-drying, as the characteristic ketone peak of camphor (1743 cmD¹) was absent while PCL signatures were preserved (Figure 1J). Through-thickness permeability and surface wettability were further verified by rapid wicking of aqueous dye across the membrane (Figure 1K), consistent with an open, hydrophilic, and vertically interconnected pore network suitable for sustained ALI culture.

Mechanical characterization demonstrated tunable compliance linked to pore architecture and geometry. Increasing porosity reduced the elastic modulus from 23.09 MPa for nonporous PCL to 8.87 MPa for the porous membrane, while preserving elastic behavior within the applied cyclic strain range (Figure 1L,M; Figure S5; Table S3). Incorporation of the hemispherical dome geometry increased tensile strain at break without substantially altering modulus (Figure S6), indicating enhanced deformability arising from three-dimensional curvature. Subsequent lipase-mediated pore enlargement further reduced modulus to 5.34 MPa, approaching reported values for native lung tissue [53]. After 14 days of ALI culture under cyclic deformation, no significant loss of mechanical integrity was observed, confirming structural stability under physiologic loading.

Together, the high degree of pore interconnectivity, ultrathin dome regions, and three-dimensional alveolar-shaped array establish a structurally defined membrane architecture that balances minimized diffusion resistance with mechanical robustness. Tunable porosity and geometry enable compliance approaching that of native lung tissue while preserving stability under prolonged cyclic deformation. This combination of interconnectivity, controlled thickness, and curvature-defined strain provides a transport-competent scaffold designed to support physiologic epithelial-endothelial interfaces and enable quantitative interrogation of cross-barrier mass transfer.

### 2.2 Transport performance and functional barrier integration of the interstitium-mimicking membrane

To evaluate transport and barrier behavior under physiologically relevant conditions, the interstitium-mimicking membrane was integrated between apical and basolateral PDMS chambers to construct a compartmentalized alveolar interface (Figure 2A; Figure S7A). This platform was designed primarily to evaluate how the membrane architecture influences transport, barrier formation, and cellular responses under physiologically relevant conditions. In this configuration, epithelial cells are cultured on the apical membrane surface under air–liquid interface (ALI) conditions, while endothelial cells are cultured on the basolateral side, recreating the epithelial–endothelial arrangement of the native air–blood barrier. The apical chamber permits direct aerosol or gas exposure, and the basolateral chamber supports continuous perfusion. Cyclic deformation is applied through programmable actuation of the apical ceiling, transmitting controlled strain to the dome-shaped membrane architecture. *In vivo*, alveoli experience ∼4% linear strain during quiet breathing and up to ∼12% during exercise at frequencies approaching ∼0.5 Hz [54]. To remain within this physiologic-exercise range while eliciting measurable transport effects, cyclic deformation of 10% linear strain at 0.33 Hz was applied throughout transport and exposure experiments. For scalable experimentation, the system was implemented in eight-unit array formats compatible with standard 12-well plates (Figure S7B-D). The array enables parallel studies in a compact footprint, while allowing each device to be configured with distinct membrane geometries/porosities and independently programmed cyclic deformation. Modular layout therefore supports space-efficient, high-throughput experimentation with flexible condition multiplexing.

**Figure 2.**
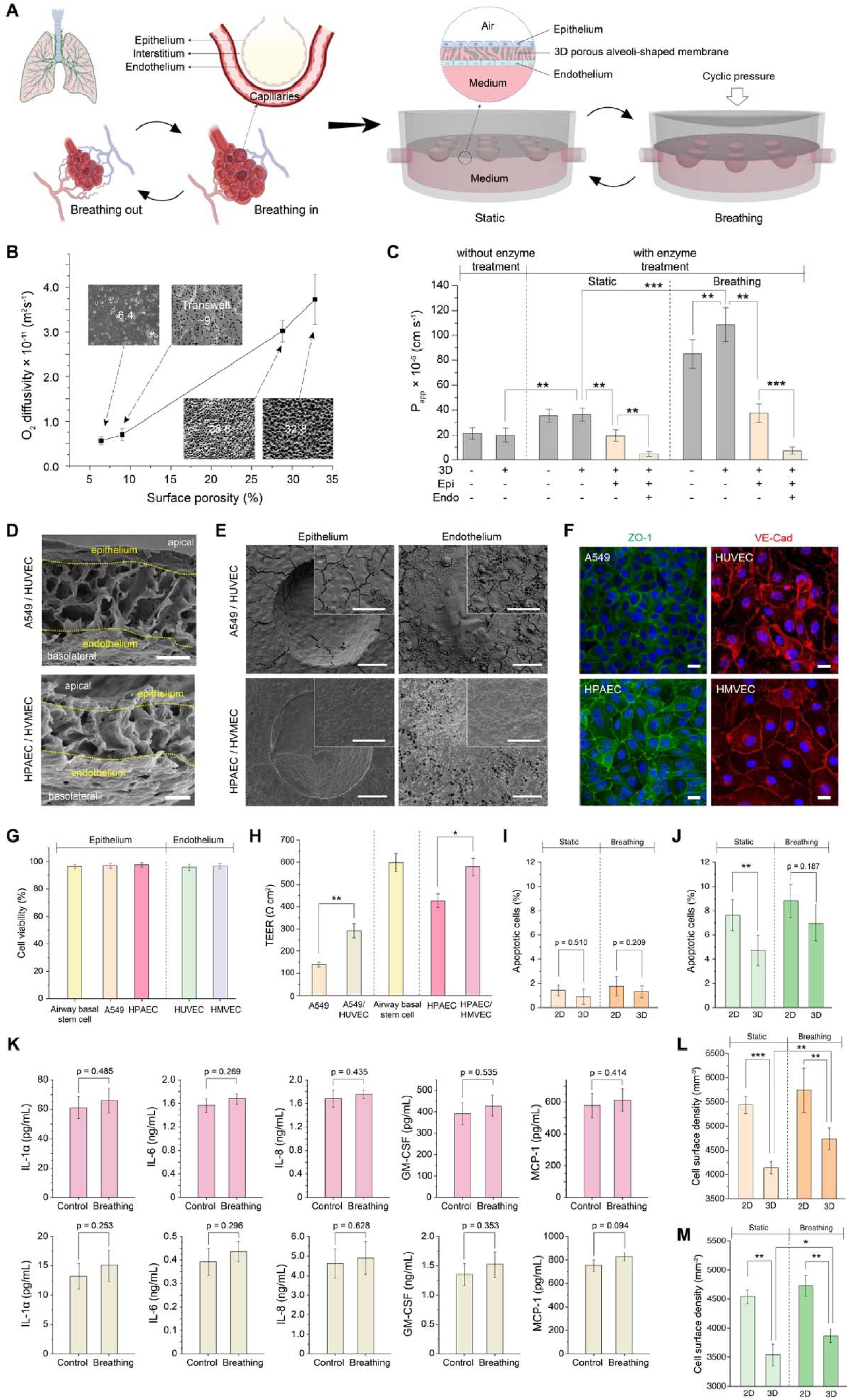
Transport performance and functional barrier integration of the interstitium-mimicking membrane under cyclic deformation. (**A**) Overview of the lung alveoli-on-chip platform. Illustration of the native alveolar-capillary structure during breathing (left) alongside the architecture of the compartmentalized microfluidic device (right). (**B**) Oxygen diffusivity of interstitium-mimicking membranes with varying porosity (6.4%, 28.8%, 32.8%) compared with a Transwell® membrane (porosity: 9.1%). Oxygen diffusivity increased with increasing membrane porosity, showing a strong positive linear correlation (R² = 0.959, p < 0.0001) (n = 4). (**C**) Apparent permeability (P_app_) of 4 kDa dextran through porous PCL membranes and epithelial-endothelial co-cultures (A549-HUVEC) under static and cyclic deformation conditions. Alveolus-shaped (3D) membranes were compared with flat (2D) membranes. (n = 5). (**D**) Representative cross-sectional SEM images of epithelial-endothelial co-cultures on the alveolus-shaped porous PCL membrane: A549-HUVEC (left) and HPAEC-HMVEC (right) under ALI with cyclic deformation. Scale bar: 5 µm. (**E**) Representative top-view SEM images of the same co-culture configurations. Scale bar: 50 µm. (**F**) Representative immunostaining of epithelial (ZO-1) and endothelial (VE-cadherin) junctions in A549-HUVEC (top) and HPAEC-HMVEC (bottom) co-cultures. Scale bars: 20 µm. (**G**) Quantification of cell viability under cyclic deformation for A549–HUVEC (8 days), HPAEC–HMVEC (10 days), and airway basal stem cell cultures (3 weeks). (**H**) TEER of the three culture configurations measured under static conditions after barrier formation (n = 4). (**I,J**) Quantification of apoptosis in A549 cells (I) and HUVECs (J) under static and cyclic deformation conditions, assessed in flat (2D) and alveolus-shaped (3D) regions (n = 5). (**K**) Secretion levels of IL-1α, IL-6, IL-8, GM-CSF, and MCP-1 under static and cyclic deformation conditions in HPAEC–HMVEC (top) and A549–HUVEC (bottom) co-cultures (n = 4). (**L,M**) Surface cell density of A549 cells (L) and HUVECs (M) under static and cyclic deformation conditions, comparing flat (2D) and alveolus-shaped (3D) regions (n = 5).

The membrane was evaluated across multiple epithelial-endothelial configurations to assess generalizability. Two primary co-culture systems were established: A549 epithelial cells with HUVECs and human primary alveolar epithelial cells (HPAECs) with human lung microvascular endothelial cells (HMVECs). The primary epithelial cells showed negligible expression of airway basal marker P63 (Figure S8), confirming alveolar phenotype. In addition, airway basal stem cells were cultured under ALI conditions, given reports of their migration into distal lungs during injury and repair [55, 56]. All configurations formed confluent monolayers on the porous scaffold under cyclic deformation.

Intrinsic transport properties were first assessed under acellular conditions. Oxygen diffusivity measurements demonstrated that the interstitium-mimicking membrane exhibited approximately 5.3-fold higher oxygen transport than conventional Transwell® membrane (Figure 2B; Table S4) [57, 58], consistent with its high open fraction and vertically interconnected pore network. Similarly, permeability to 4 kDa FITC-dextran increased by 1.83-fold following lipase-mediated pore enlargement, confirming that architectural tuning directly reduces membrane-imposed mass transfer resistance (Figure 2C). Under cyclic deformation, cell-free interstitium-mimicking membranes displayed 1.27-fold increased permeability relative to flat membranes, reflecting strain-induced modulation of effective pore geometry and curvature-dependent deformation, consistent with finite element simulation (Figure S9). These results establish the membrane as a low-resistance, strain-responsive transport scaffold. Unlike conventional flat-membrane lung-on-chip systems that generate strain through lateral vacuum actuation [1, 19, 59], the present platform applies normal pressure loading to a hemispherical membrane, producing curvature-dependent and spatially heterogeneous local strain fields that are difficult to reproduce in flat-membrane systems.

The ability of this transport-competent membrane to support formation of a functional epithelial-endothelial barrier was next evaluated. Surface wettability was adjusted using extracellular matrix coatings to promote uniform cell attachment (Figure S10). Under ALI conditions, epithelial and endothelial cells formed confluent monolayers on opposing membrane surfaces, as confirmed by cross-sectional SEM imaging (Figure 2D,E; Figures S10,S11). No evidence of epithelial cell penetration through the membrane or invasion into the endothelial compartment was observed. Continuous intercellular junctions were verified by ZO-1 staining in epithelial layers and VE-cadherin staining in endothelial layers (Figure 2F; Figure S13). Cell viability exceeded 95% under cyclic deformation (Figure 2G). Transepithelial/transendothelial electrical resistance (TEER) of epithelial monolayers reached values within reported ranges for functional alveolar barrier models [19]. TEER values increased further upon epithelial-endothelial co-culture (Figure 2H; Figure S14), indicating establishment of a more stable, less permeable and strain-tolerant interface [60]. Notably, barrier properties varied by epithelial phenotype. Airway basal stem cell cultures exhibited elevated TEER values even in the absence of endothelial co-culture, indicating formation of a particularly tight epithelial barrier.

To evaluate the influence of cyclic deformation and three-dimensional curvature on baseline cell behavior, epithelial-endothelial co-cultures were examined under physiologic strain. Under 10% linear strain at 0.33 Hz, the membrane maintained structural integrity (Figure 1L), and confluent monolayers remained firmly adhered on both surfaces (Figure 2D-F). Apoptosis, assessed by cleaved caspase-3 staining, showed no significant increase under cyclic deformation in either epithelial or endothelial layers (Figure 2I,J; Figure S15), indicating that applied strain alone did not induce overt cytotoxic stress. Baseline cytokine profiling under strain revealed only modest, non-significant changes in IL-1α, IL-6, IL-8, GM-CSF, and MCP-1 (Figure 2K), confirming that cyclic deformation within the applied range does not independently provoke inflammatory activation.

Since the epithelial layer serves as a protective interface while the capillary endothelium governs fluid transport within the pulmonary microenvironment [61, 62], transport measurements were then repeated in the presence of intact epithelial-endothelial layers. As expected, oxygen diffusivity and dextran permeability decreased relative to acellular conditions due to barrier formation (Figure 2C; Figure S16,S17). Cyclic deformation further increased cross-barrier permeability 2.97-fold relative to static culture, indicating strain-induced pore opening, in-plane cell and membrane deformation, and cyclic pressure promoting downward particle movement.

To isolate curvature-dependent effects, planar and dome-shaped membranes were compared. A549 and HUVEC monolayers on curved alveolar surfaces exhibited reduced surface cell density relative to planar substrates (Figure 2L,M), whereas HPAEC–HMVEC co-cultures showed no significant geometry-dependent differences (Figure S18). Cyclic strain increased surface coverage on dome-shaped substrates but had minimal effect on planar controls, consistent with curvature-mediated mechanical cues influencing cellular organization [63–65].

Collectively, these findings demonstrate that the interstitium-mimicking membrane combines high intrinsic permeability with the capacity to support a functional, mechanically robust alveolar barrier. The architectural advantages conferred by high interconnectivity, minimized diffusion distance, and dome-defined deformation are retained after epithelial-endothelial integration, establishing a transport-competent platform for quantitative interrogation of cross-interface mass transfer.

### 2.3 Quantitative aerosol exposure reveals dose-dependent alveolar barrier responses

With a transport-competent epithelial-endothelial interface established, controlled aerosol deposition was performed under ALI conditions to interrogate barrier responses to carbonaceous particulate exposure (Figure 3A) [66]. The particulate used in this study was a well-characterized carbon-dominant coal dust (>95% carbon by composition) enriched in nanoscale fractions and polycyclic aromatic hydrocarbons (PAHs) [42]. Such physicochemical features are characteristic of combustion-derived and certain mining-associated aerosols. Particles were delivered directly to the apical surface at defined surface mass densities, enabling quantitative and reproducible dosing while maintaining basolateral perfusion and cyclic mechanical loading. Dry particles with a primary size of ∼0.4 µm [42] formed aerosols with a median aerodynamic diameter of approximately 1.0 µm (Figure 3B), placing them within the respirable size range relevant to distal lung exposure.

**Figure 3.**
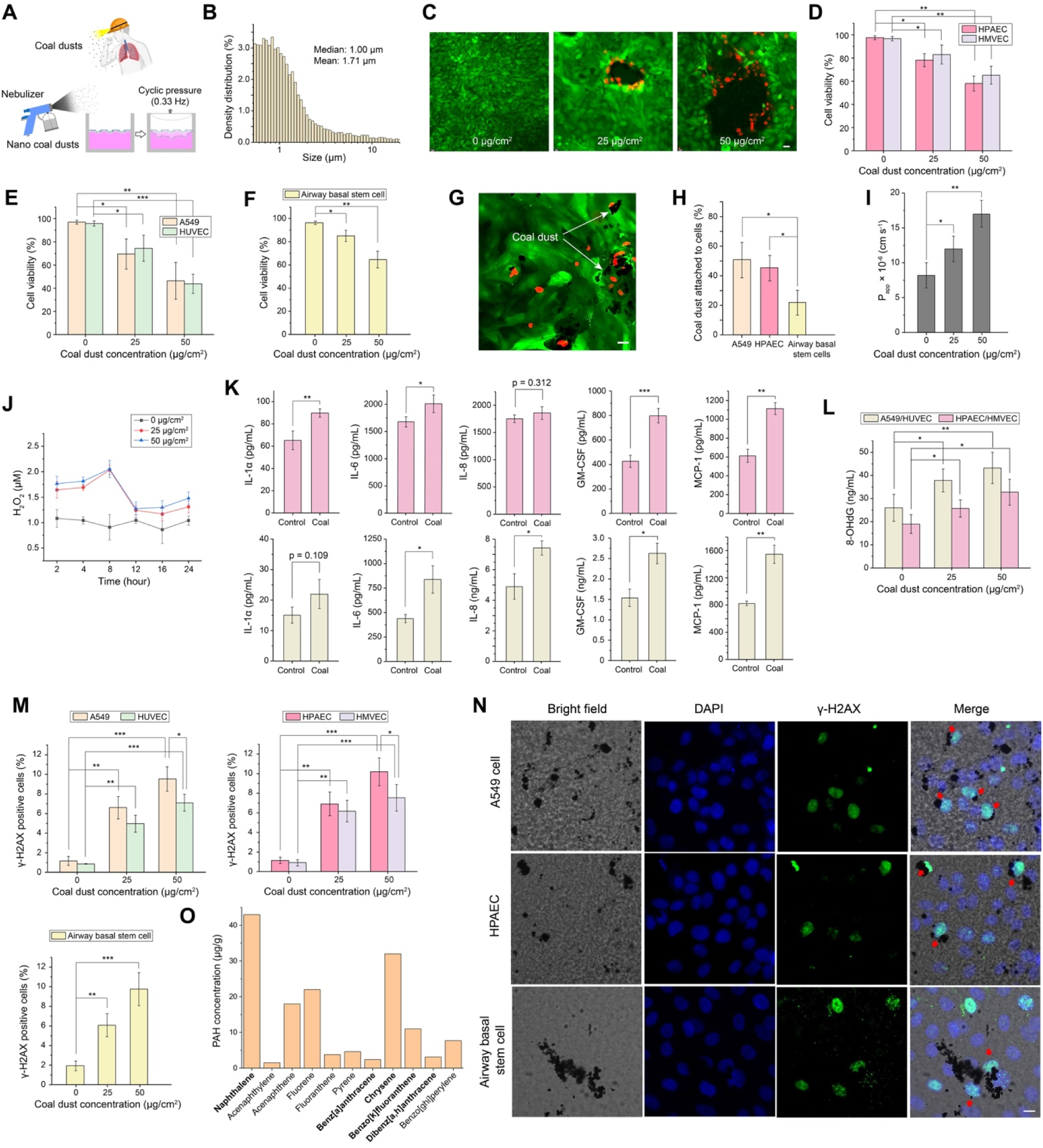
Carbon-dominant coal dust induces dose-dependent alveolar barrier responses. (**A**) Schematic of the aerosol exposure protocol in the alveoli-on-chip system. Under ALI conditions, carbon-dominant coal dust was delivered to the apical chamber using a nebulizer, followed by 24 hours of cyclic deformation (10% linear strain, 0.33 Hz). (**B**) Aerodynamic particle size distribution of aerosolized coal dust, measured using an Aerodynamic Particle Sizer (TSI 3321), showing a mean diameter of 1.71 µm and a median of 0.99 µm. (**C**) Representative fluorescence images showing concentration-dependent disruption of HPAEC epithelial layers following coal dust exposure. (**D-F**) Quantification of cell viability in HPAEC–HMVEC co-cultures, A549–HUVEC co-cultures, and airway basal stem cell monocultures after exposure to coal dust at surface loadings of 0, 25, and 50 µg cmD² (n = 4). (**G**) Representative live (green)/dead (red) staining of airway basal stem cells following 50 µg cmD² coal dust exposure. Scale bar: 20 µm. (**H**) Percentage of coal dust attached to epithelial cell surfaces 24 hours after exposure in airway basal stem cells, A549 cells, and HPAECs. (**I**) Apparent permeability (P_app_) to 4 kDa FITC-dextran across A549–HUVEC co-cultures after 24 hours of coal dust exposure under ALI with cyclic deformation (0, 25, and 50 µg cmD²; n = 5). (**J**) ROS generation, quantified by hydrogen peroxide (HDOD) concentration in A549-HUVEC co-cultures at 2-24 hours following coal dust exposure (0, 25, and 50 µg cmD²; n = 5). (**K**) Secretion of IL-1α, IL-6, IL-8, GM-CSF, and MCP-1 in HPAEC–HMVEC (top) and A549-HUVEC (bottom) co-cultures 24 hours after 25 µg cmD² coal dust exposure under ALI with cyclic deformation (n = 4). (**L**) Oxidative DNA damage measured by 8-OHdG ELISA in co-culture systems 24 hours after exposure to coal dust (0, 25, and 50 µg cmD²; n = 5). (**M**) Percentage of γ-H2AX-positive cells in A549-HUVEC, HPAEC-HMVEC, and airway basal stem cell cultures after coal dust exposure (0, 25, and 50 µg cmD²; n = 4). (**N**) Representative immunostaining of γ-H2AX indicating DNA double-strand breaks in epithelial cell types following 24 hours exposure to 50 µg cmD² coal dust. Scale bar: 10 µm. (**O**) Quantified concentrations of PAHs detected in the coal sample. Compounds shown in bold are classified by the International Agency for Research on Cancer (IARC) as Group 1 or Group 2 carcinogens.

Carbonaceous particulate exposure produced dose-dependent reductions in epithelial-endothelial viability across all three culture configurations. In HPAEC-HMVEC co-cultures, viability declined to approximately 78% (epithelium) and 83% (endothelium) at 25 µg cmD² and further to ∼58% and ∼65% at 50 µg cmD², respectively (Figure 3C,D; Figure S19). Similar dose-dependent trends were observed in A549-HUVEC and airway basal stem cell cultures (Figure 3E-G), indicating that particulate-induced injury is not restricted to a specific epithelial phenotype. Comparable reductions in epithelial and endothelial viability suggest that injury propagates across the integrated interface rather than remaining confined to the apical layer.

Particle-cell interactions differed among epithelial phenotypes. In differentiated mucociliary airway basal stem cell cultures, particulate attachment was markedly reduced compared with non-ciliated epithelial models (Figure 3H; Figure S20), consistent with enhanced mucociliary clearance mechanisms [67]. In contrast, alveolar epithelial layers exhibited greater surface retention of particulates, reflecting phenotype-dependent exposure dynamics.

Barrier function deteriorated in parallel with viability loss. Apparent permeability to 4 kDa FITC-dextran increased with exposure dose under ALI conditions with cyclic strain (Figure 3I; Figure S21), consistent with concentration-dependent disruption of epithelial monolayers and junctional compromise under particulate stress [62, 68, 69].

Recovery capacity was strongly dose-dependent. At 50 µg cmD², primary epithelial and endothelial layers exhibited sustained viability loss through day 5, indicating severe and persistent injury. At 25 µg cmD² or lower, partial restoration of viability was observed by day 5 (Figure S22), suggesting activation of repair or adaptive responses following sublethal exposure. Notably, primary epithelial cells cultured on Transwell® membrane failed to exhibit comparable recovery at equivalent dose (Figure S23), indicating that dynamic mechanical stimulation and epithelial-endothelial coupling in the membrane platform influence injury resolution.

Oxidative stress responses were quantified to assess early molecular injury following particulate exposure. Reactive oxygen species (ROS) generation, quantified by hydrogen peroxide release, increased in a dose-dependent manner and peaked within 8-hour post-exposure across co-culture configurations (Figure 3J), indicating rapid redox imbalance induced by carbonaceous particulates [70]. Pro-inflammatory cytokines, including IL-6, GM-CSF, and MCP-1, were significantly elevated under 25 µg cmD² exposure in both immortalized and primary epithelial–endothelial systems (Figure 3K), reflecting coordinated intercellular signaling across the barrier interface. Markers of genomic injury further revealed dose-dependent accumulation of oxidative DNA damage and double-strand breaks. Levels of 8-hydroxy-deoxyguanosine increased with exposure (Figure 3L), and γ-H2AX–positive nuclei were elevated in epithelial and endothelial layers at higher particulate loads (Figure 3M,N). Chemical analysis identified multiple polycyclic aromatic hydrocarbons (PAHs) within the particulate fraction (Figure 3O), compounds known to undergo metabolic activation and generate oxidative DNA lesions and replication-associated double-strand breaks. The presence of these PAHs is consistent with the observed genotoxic signatures [71, 72]. These findings demonstrate that controlled aerosol deposition on the interstitium-mimicking membrane induces reproducible oxidative, inflammatory, and genotoxic responses under physiologically relevant mechanical conditions [73, 74].

### 2.4 Matched cumulative dose reveals divergent acute-chronic injury response programs

To distinguish the influence of exposure pattern from total dose, matched cumulative dosing regimens were compared under ALI conditions (Figure 4A). A single 25 µg cmD² exposure defined the acute condition, whereas five sequential 5 µg cmD² exposures over 17 days defined the chronic condition, yielding identical total surface mass deposition.

**Figure 4.**
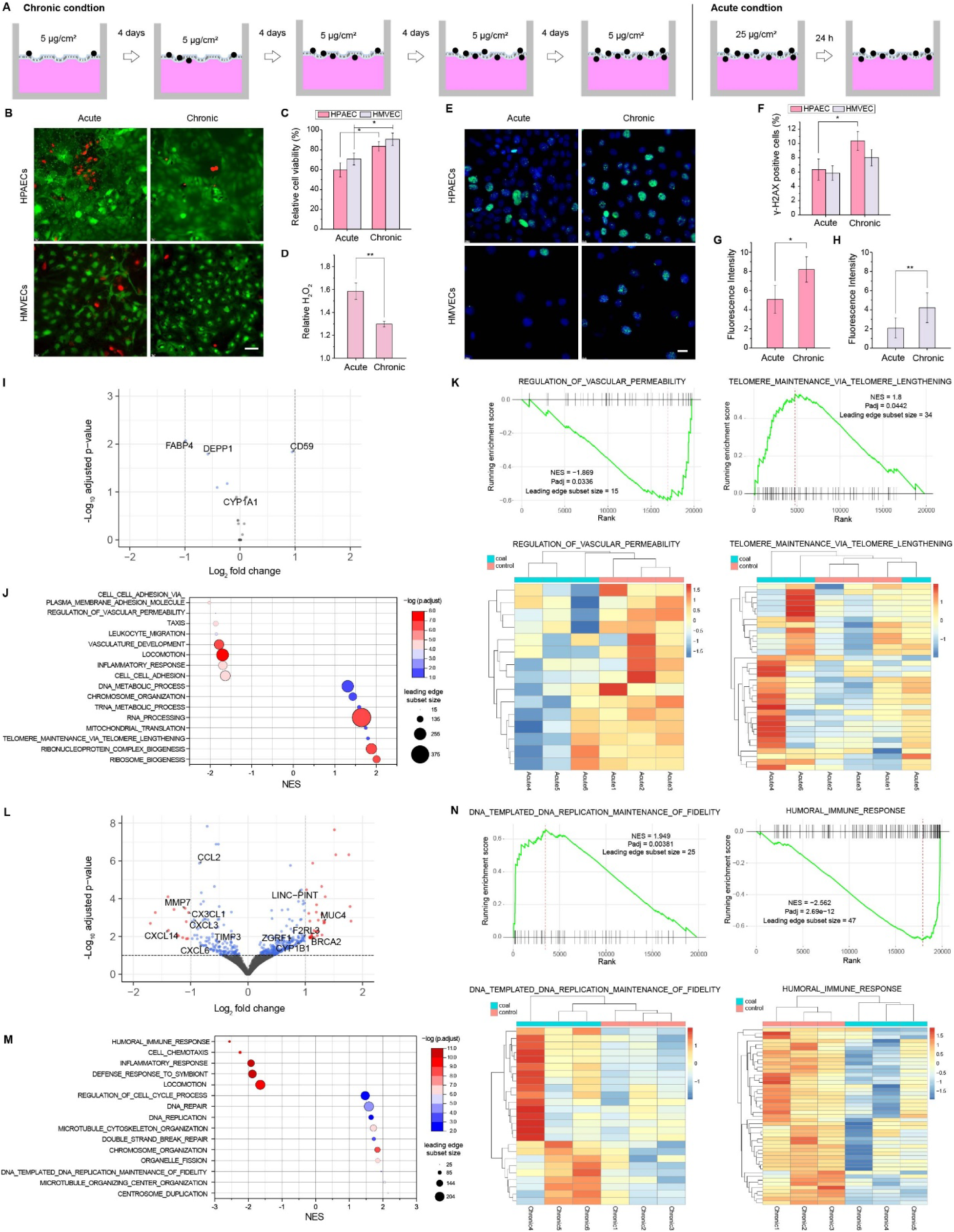
Matched cumulative coal dust exposure reveals divergent acute and chronic injury programs. (**A**) Experimental design for matched cumulative dose comparison. In both regimens, a total surface loading of 25 µg cmD² coal dust was applied under ALI conditions. The acute condition consisted of a single 25 µg cmD² exposure with analysis at 24 hours, whereas the chronic condition consisted of five sequential 5 µg cmD² exposures over 17 days prior to analysis. (**B**) Representative live/dead fluorescence images of epithelial and endothelial layers following acute and chronic exposure. Scale bar: 50 µm. (**C**) Relative cell viability after acute and chronic exposure. Values are normalized to corresponding untreated controls (n = 4). (**D**) Relative HDOD levels following acute and chronic exposure, normalized to control levels (n = 4). (**E**) γ-H2AX immunofluorescence staining revealing DNA double-strand damage in epithelial and endothelial cells after acute and chronic exposures. Scale bar: 20 µm. (**F**) Relative percentage of γ-H2AX–positive cells following acute and chronic exposure, normalized to control levels (n = 4). (**G,H**) Quantification of γ-H2AX fluorescence intensity in HPAECs (G) and HMVECs (H) under acute and chronic exposure. Fluorescence intensity was normalized to cell area (n = 4). (**I**) Volcano plot of differentially expressed genes (DEGs) under the acute condition (n = 3). (**J**) Gene set enrichment analysis (GSEA) dot plot of significantly enriched Gene Ontology Biological Process (GOBP) terms under the acute condition. (**K**) Representative GSEA enrichment plots and leading-edge gene heatmaps for acute exposure compared with controls. (**L**) Volcano plot of DEGs under the chronic condition (n = 3). (**M**) GSEA dot plot of significantly enriched GOBP terms under the chronic condition. (**N**) Representative GSEA enrichment plots and leading-edge gene heatmaps for chronic exposure compared with controls.

Despite equivalent cumulative dose, biological outcomes diverged markedly. Acute exposure induced substantial reductions in epithelial and endothelial viability accompanied by elevated ROS levels within 24 hours (Figure 4B-D). In contrast, chronic low-dose exposure preserved overall viability and exhibited lower peak ROS generation at matched time points. These findings indicate that high instantaneous particulate burden drives rapid cytotoxic stress, whereas repeated low-level exposure permits partial cellular adaptation. Genotoxic outcomes, however, followed an opposing trend. Chronic exposure produced higher fractions of γ-H2AX-positive nuclei and greater fluorescence intensity compared with the acute regimen (Figure 4E-H), consistent with sustained DNA damage accumulation despite preserved short-term viability. This likely reflects the cumulative nature of genotoxic stress and the incomplete resolution of damage arising from sustained exposure over time, despite a lower instantaneous oxidative burden [75].

To delineate the molecular programs underlying exposure-pattern divergence, bulk RNA sequencing was performed on epithelial-endothelial co-cultures following acute and chronic regimens. Under acute exposure (25 µg cmD², 24 h), differentially expressed genes reflected an immediate cytotoxic stress response. FABP4 and DEPP1, associated with lipid metabolism and stress-regulated autophagy, were downregulated, whereas CD59, a complement regulatory protein, was upregulated (Figure 4I) [76–78]. Although CYP1A1, a canonical xenobiotic-metabolizing enzyme, exhibited an upward trend, its induction did not reach statistical significance (Figure S24) [79]. Gene set enrichment analysis (GSEA) revealed prominent activation of ribosome biogenesis, RNA processing, and mitochondrial translation pathways (Figure 4J,K; Figure S25; Table S5), accompanied by reduced cell–cell adhesion and inflammatory pathway signatures. These features are consistent with a regenerative biosynthesis program emerging from a surviving cell population following acute oxidative injury. At 24 hours, transcriptomic profiles likely reflect cells that withstood the initial ROS surge and barrier disruption observed in earlier measurements, engaging compensatory metabolic and chromatin remodeling processes. In contrast, chronic exposure (five sequential 5 µg cmD² doses) elicited a transcriptional program dominated by genome stability and DNA repair pathways. Genes linked to environmental pollutant exposure, including F2RL3 and CYP1B1, were markedly upregulated (Figure 4L; Figure S26) [80, 81]. Concomitantly, DNA damage response and repair-associated genes such as BRCA2 and ZGRF1, as well as tumor-suppressive noncoding transcripts including LINC-PINT, were elevated [82–85]. GSEA demonstrated enrichment of double-strand break repair, DNA replication, chromosome organization, centrosome duplication, and microtubule organizing center pathways (Figure 4M,N; Figure S27; Table S5), indicating sustained genomic maintenance activity. In parallel, multiple chemokine and inflammatory mediators (e.g., CCL2, CXCL3, CXCL6, CXCL14, CX3CL1) and matrix remodeling genes (MMP7, TIMP3) were downregulated, suggesting attenuation of acute inflammatory signaling in favor of genomic preservation [86–88].

Notably, DNA damage-associated pathways diverged sharply between regimens. Acute exposure showed limited enrichment of DNA-related processes beyond telomere maintenance, consistent with immediate oxidative insult followed by survival-driven recovery. Chronic exposure, however, produced broad activation of replication-coupled repair and chromosome stability programs, aligning with persistent γ-H2AX elevation despite preserved short-term viability. Together, these transcriptomic distinctions indicate that high-intensity acute exposure favors rapid cytotoxic collapse and compensatory biosynthesis in surviving cells, whereas repeated low-dose exposure promotes sustained genomic stress and repair without overt apoptosis. These findings demonstrate that the interstitium-mimicking membrane platform resolves exposure-pattern-dependent molecular programs that are not predictable from cumulative dose alone.

### 2.5 Therapeutic intervention attenuates particulate-induced injury

To demonstrate translational applicability, therapeutic interventions targeting oxidative and inflammatory pathways were evaluated during particulate exposure. Antioxidant treatment with vitamins C and E was administered via the basolateral compartment during apical aerosol challenge [89]. Co-treatment improved epithelial and endothelial viability in a dose-dependent manner (Figure 5A) and reduced hydrogen peroxide levels (Figure 5B), indicating attenuation of oxidative injury.

**Figure 5.**
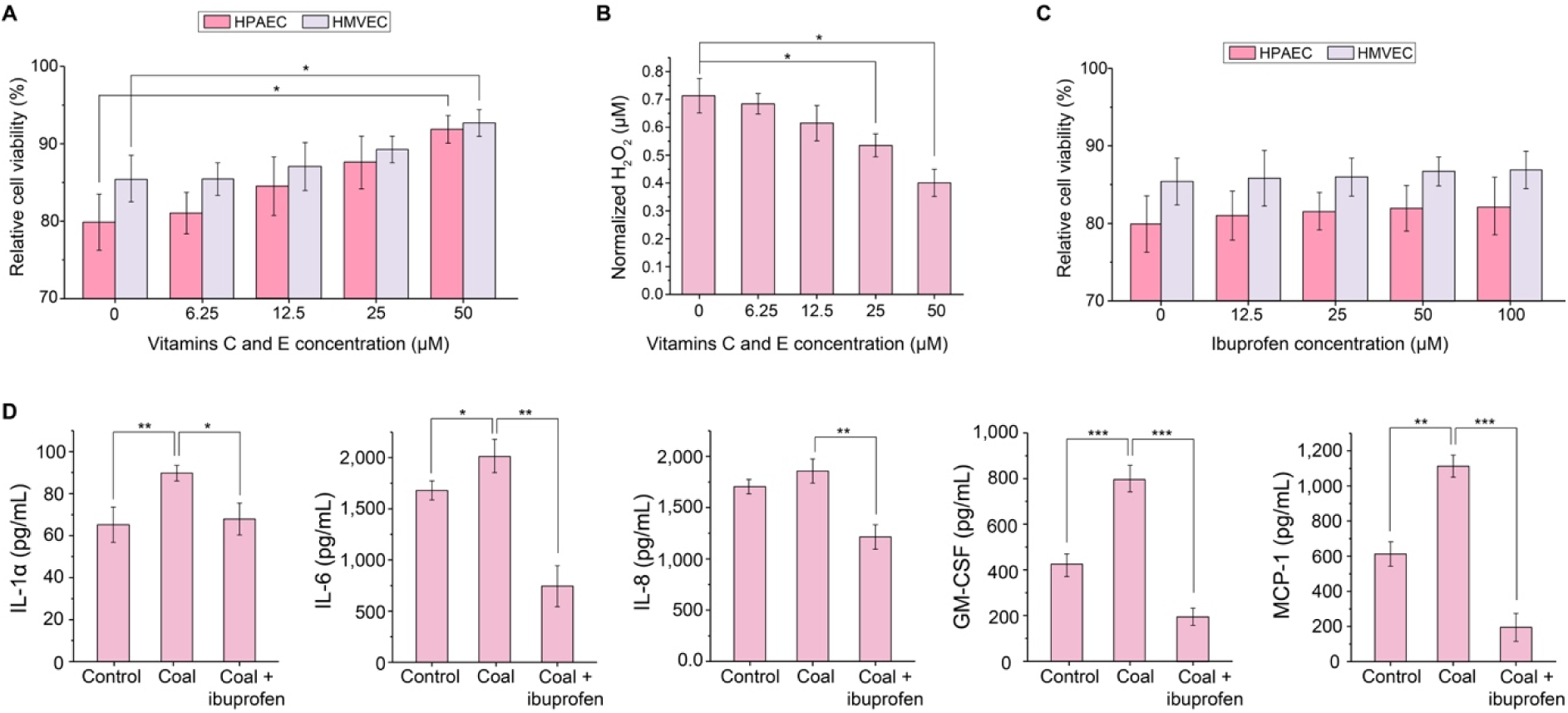
Therapeutic intervention attenuates coal dust-induced alveolar injury. (**A**) Quantification of relative viability in HPAEC–HMVEC co-cultures 24 h after administration of vitamins C and E (0–50 µM) to the basolateral channel under coal dust challenge (25 µg cmD²) (n = 4). (**B**) HDOD levels measured 24 h after co-treatment with vitamins C and E during coal dust exposure (25 µg cmD²) (n = 4). (**C**) Relative cell viability of HPAEC-HMVEC co-cultures measured 24 hours after simultaneous administration of ibuprofen (0–100 µM) to the basolateral chamber during coal dust exposure (25 µg cmD²). Values are normalized to untreated controls (n = 4). (**D**) Cytokine secretion (IL-6, GM-CSF, MCP-1) measured in HPAEC-HMVEC co-cultures under three conditions: control (ALI with cyclic deformation), coal dust exposure (25 µg cmD²), and coal dust plus ibuprofen (100 µM). Basolateral media were collected 24 hours after exposure (n = 4).

The anti-inflammatory agent ibuprofen was similarly assessed for its ability to mitigate cytokine responses [90]. Ibuprofen did not significantly alter overall epithelial or endothelial viability but significantly reduced secretion of IL-6, GM-CSF, and MCP-1 following particulate exposure (Figure 5C,5D), indicating selective suppression of inflammatory signaling without reversal of cytotoxic injury.

These findings highlight the capacity of the membrane-enabled alveolar interface to evaluate therapeutic strategies under controlled, physiologically relevant aerosol exposure. The platform therefore provides a structurally defined system for preclinical assessment of interventions targeting inhalation-driven lung injury.

## 3. Conclusion and Discussion

This work introduces an interstitium-mimicking porous membrane engineered to replicate key structural and mechanical features of the alveolar septum. Serving as the structural and functional core of an alveoli-on-chip platform, the membrane combines high pore interconnectivity, reduced diffusion distance, and mechanical compliance within a three-dimensional alveolar geometry. Integration of this membrane into a compartmentalized epithelial-endothelial system enables quantitative investigation of transport, barrier, and aerosol-exposure dynamics under physiologically relevant mechanical conditions. Fabricated via a dual-templated nonsolvent-induced phase separation strategy combined with controlled enzymatic pore enlargement, the membrane achieves high overall porosity (∼40%) with 97% pore interconnectivity while incorporating a three-dimensional alveolar dome array (∼200 µm units) containing locally thinned regions (∼2.5 µm). This architecture simultaneously minimizes diffusion distance, preserves vertical transport continuity, and distributes mechanical strain through curvature-defined deformation fields. The resulting structure reduces membrane-imposed transport resistance while maintaining compliance and stability under cyclic loading, establishing a transport-competent scaffold that approximates native septal geometry.

Integrated between apical and basolateral chambers, the interstitium-mimicking membrane forms the structural core of a compartmentalized alveolar interface that supports air-liquid culture, perfusion, and programmable mechanical deformation. The dome-shaped geometry introduces curvature-dependent strain heterogeneity that modulates mass transfer across the epithelial-endothelial interface while preserving barrier integrity. Compared with conventional flat, low-porosity thick membranes, this design maintains enhanced oxygen transport and strain-responsive permeability even after epithelial-endothelial integration. By restoring physiologically relevant exchange pathways within a mechanically dynamic compartmental system, the membrane enables quantitative interrogation of cross-interface transport and barrier behavior under controlled aerosol exposure.

The platform further resolves multidimensional biological responses to aerosol exposure. Controlled carbonaceous particulate deposition induces reproducible oxidative, inflammatory, and genotoxic injury programs, while matched cumulative dosing reveals that exposure pattern, not total dose alone, governs divergent acute and chronic cellular outcomes. Acute high-intensity exposure favors rapid cytotoxic stress and compensatory biosynthesis in surviving cells, whereas repeated low-dose exposure promotes sustained DNA repair and genome-maintenance programs despite preserved short-term viability. Compartment-resolved experiments demonstrate that severe apical exposure can propagate downstream vascular consequences, highlighting the ability of the membrane-based system to dissect spatial aspects of injury propagation. Therapeutic interventions targeting oxidative and inflammatory pathways further demonstrate the utility of the platform for translational evaluation under physiologically relevant aerosol conditions.

Importantly, the interstitium-mimicking membrane provides a structurally defined framework for interrogating damage and repair across alveolar diseases in which epithelial-endothelial exchange and mechanical stress are central drivers of pathology. Conditions such as viral and bacterial pneumonia, COPD, ARDS, pulmonary fibrosis, asthma, ventilation-induced injury, smoking-related diseases, and lung cancer involve disruption of barrier integrity, altered transport dynamics, and mechanically mediated remodeling. By restoring physiologically relevant diffusion distance, interconnectivity, and strain responsiveness within a compartmentalized system, the membrane platform enables mechanistic dissection of how geometry, transport, and exposure pattern influence disease progression and therapeutic response.

Beyond pulmonary applications, the same architectural principles—high pore interconnectivity, minimized diffusion distance, and mechanically compliant three-dimensional curvature—provide a generalizable strategy for constructing compartmentalized microphysiological systems that require controlled bidirectional exchange of soluble factors, extracellular vesicles, or particulate matter while preserving distinct cellular domains. Such design may be extended to other barrier tissues, including gut, blood–brain barrier, placental, and tumor–stromal interfaces. By unifying geometric fidelity, transport competence, and mechanical coupling within a scalable fabrication framework, the interstitium-mimicking membrane advances lung-on-chip technology from structural mimicry toward functional replication of barrier transport dynamics, establishing a broadly adaptable materials platform for studying injury, repair, and therapeutic intervention in compartmentalized tissue systems.

## 4. Experimental Section

### 4.1 Fabrication of alveolar-shaped mold

The SU-8 mold used for membrane fabrication was prepared on a 4-inch silicon wafer. Prior to photoresist coating, the wafer was treated with UV/ozone for 10 min (Samco UV-1 Ozone Cleaner) and dehydrated at 200 °C for an additional 10 min. SU8-2075 (MicroChem) was subsequently spin-coated to obtain a photoresist layer approximately 60 µm thick. After a soft-bake step at 65 °C for 2 min and 95 °C for 8 min, the wafer was exposed to 365 nm ultraviolet light (5 mW cm^-2^, 60 s) using a Karl Suss MA56 mask aligner. Post-exposure baking was performed at 65 °C for 1.5 min followed by 95 °C for 6.5 min. The patterned structures were developed in SU-8 developer for 6 min and sequentially rinsed with acetone, isopropyl alcohol, and deionized water. A final hard-bake at 95 °C for 15 min was carried out to enhance adhesion between the SU-8 features and the substrate. Mold thickness was verified using a KLA Tencor P-15 profilometer.

### 4.2 Interstitium-mimicking porous PCL membrane fabrication

A 10 wt% PCL solution was obtained by dissolving PCL pellets (Mn = 80,000, Sigma-Aldrich) in acetone under continuous stirring at 60 °C for 3 h. Camphor (150 wt% with respect to PCL; 96%, Sigma-Aldrich) was then introduced as a porogen and allowed to dissolve completely during an additional 1 h mixing period. After equilibration to room temperature, the polymer mixture was deposited onto a patterned SU-8 mold by spin coating (400 rpm, 5 s). Membrane solidification was achieved by immersing the coated mold in distilled water for 1 h. Subsequently, camphor was removed via freeze-drying for 3 h, yielding a porous membrane structure. Enzymatic degradation studies were performed by incubating the membranes in lipase-containing PBS at 37 °C.

### 4.3 Membrane Characterization

The morphology and pore architecture of the fabricated PCL membranes were examined using scanning electron microscopy (SEM; Quanta 600 FEG, FEI, USA). To visualize the internal structure, membranes were cryofractured in liquid nitrogen prior to imaging. For cell-seeded samples, fixation was performed with 2.5% glutaraldehyde for 1 h at room temperature, followed by three PBS washes. Samples were then treated with 1% osmium tetroxide for 1 h at 4 °C and dehydrated through a graded ethanol series (50–100%). Additional dehydration was carried out using hexamethyldisilazane (HMDS, Sigma-Aldrich) for 15 min before air drying. Prior to SEM analysis, all specimens were coated with a thin layer of gold. Membrane pore size and surface porosity were quantified from SEM micrographs using ImageJ software. For pore size measurements, 200 individual pores were analyzed from four representative images for each experimental group. Surface porosity was determined from the same number of images.

The overall porosity (*p*) of the porous PCL membranes was estimated from the ratio of solid volume (Vs) to total membrane volume (Vt) according to: *p* (%) = (1 - V_s_/V_t_) x 100. Total membrane volume was derived from nonporous spin-coated PCL membranes, whereas the solid volume of porous membranes was calculated from the membrane mass and PCL density (V_s_ = m/ρ_PCL_). The density of PCL was obtained from measurements of nonporous membranes. Membrane thicknesses were determined from SEM cross-sections and incorporated into the volume calculations. A total of eight samples were analyzed to calculate the mean values and standard deviations.

The chemical composition of the porous PCL membranes was characterized by Fourier transform infrared (FT-IR) spectroscopy (PerkinElmer, USA). Spectra were collected for camphor, porous PCL membranes, and pure PCL using 32 scans per sample over the wavenumber range of 600–2200 cmD¹.

Mechanical performance of the PCL membranes was evaluated by uniaxial tensile testing using an MTS Criterion® 42 system equipped with a 50 N load cell. Samples were stretched at a constant displacement rate of 10 mm/min while stress–strain data were continuously recorded. Young’s modulus was determined from the linear region of the resulting stress–strain curves. Membrane thickness values used in the analysis were obtained from cross-sectional SEM images.

Assuming that the membrane deformation approximates a partial spherical surface, the linear strain (Δ*L*) was calculated using below, where *a* is the membrane radius and *h* is the measured axial deflection during compression:

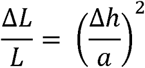

The axial deflection during compression was calculated by measuring the volume displacement of the fluid in the basolateral chamber.

### 4.4 Nano-CT

The PCL membrane was imaged using a nano-computed tomography (nano-CT) system at the Penn State Center for Quantitative Imaging (CQI). Imaging was performed at an accelerating voltage of 80 kV and power of 8 W with a beam current of approximately 100 µA using a 4× objective lens and air filter. The scan was acquired with an isotropic voxel size of approximately 1.81 µm and a field of view of approximately 3.67 mm × 3.67 mm. A total of 2401 projection images were collected over a rotation range of −180° to 10° with an exposure time of 1 s per projection. Data acquisition was performed using 1×1 detector binning (binning = 1), indicating that no pixel averaging was applied, and maximum spatial resolution was maintained during imaging. The tomographic datasets were reconstructed using the Auto CB (cone-beam) reconstruction protocol to generate cross-sectional and three-dimensional representations of the membrane microstructure. Reconstructed image stacks were analyzed in CT Analyser (CTAn, v1.13.5.1, Bruker). A total of 313 slices were analyzed using a grey-value threshold of 81∼149 to segment the solid membrane phase from the pore space. CTAn classifies pore volume connected to the exterior boundary of the analyzed volume as open porosity, whereas isolated pore volume with no exterior connection is classified as closed porosity. Pore interconnectivity was calculated as:

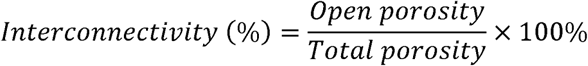

### 4.5 Oxygen diffusivity measurement

Oxygen permeability experiments were conducted by introducing 100% oxygen (1.02 atm) into the apical chamber and perfusing the basolateral channel with culture medium maintained at 37 °C (Lonza, CC-3202). Medium flow was controlled at 50 µL minD¹ using a peristaltic pump. After 5 min of oxygen exposure, dissolved oxygen levels in the medium exiting the basolateral chamber were quantified using an oxygen probe (JPB-70A, Yieryi). The resulting measurements were used to calculate oxygen diffusivity according to the equation below.

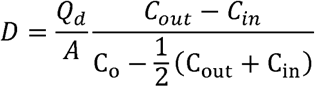

### 4.6 Chip fabrication and system construction

To fabricate the lung alveoli-on-a-chip device, an interstitium-mimicking porous PCL membrane was sandwiched between PDMS-based apical and basolateral chambers using PDMS adhesive. The assembled device was cured overnight at 45 °C to ensure bonding stability. The basolateral chamber was connected to inlet and outlet tubing for continuous medium perfusion; the outlet remained exposed to allow free drainage. A flexible PDMS lid, prepared via spin coating, was reversibly bonded on top of the apical chamber. To simulate cyclic breathing motion, a solenoid actuator (Adafruit Mini Push-pull solenoid) periodically compressed the lid at a frequency of 0.33 Hz under control of an Arduino microcontroller, inducing mechanical stretching of the porous PCL membrane. Culture medium in the basolateral chamber was perfused using a peristaltic pump (Instech P625).

### 4.7 Cell culture

Prior to cell seeding, the lung-on-chip devices were sterilized by spraying with 70% ethanol followed by 2 h of UV exposure. For experiments involving primary cells, the interstitium-mimicking membrane was pre-coated with 0.1% gelatin (Cell Biologics) at 37 °C for 5 min. Endothelial cells were first introduced into the basolateral side by inverting the device and seeding HUVECs (ATCC) or HMVECs (Lonza, passage 2) at densities of 1 × 10D and 1.5 × 10D cells, respectively. After 3 h, the device was returned to its original orientation, and epithelial cells were seeded onto the apical compartment at densities of 1 × 10D A549 cells (ATCC) or 5 × 10D HPAECs (Cell Biologics, passage 2).A549–HUVEC cocultures were maintained under liquid–liquid culture (LLC) conditions in endothelial growth medium (Lonza, CC-3162) at 37 °C and 5% CO_2_ for 7 d. ALI culture was initiated on day 7 by removing the apical medium, and cyclic membrane stretching was applied concurrently. HPAEC–HMVEC cocultures were cultured under LLC conditions using Lonza CC-3202 medium for the same duration. Following apical medium removal on day 7, the cultures were transitioned to ALI conditions and maintained for 2 d before cyclic stretching was initiated. During ALI culture, medium was continuously perfused through the basolateral compartment at 50 µL minD¹. For airway basal stem cells, the culture was performed following established protocol [91]. Briefly, the interstitium-mimicking membrane was coated with 804G-conditioned medium, followed by cell seeding at a density of 5 × 10D cells. After 24 hours of culture in expansion medium, the cells were switched to differentiation medium under ALI conditions and maintained for 3 weeks.

### 4.8 Cell viability

Live/dead staining was carried out using a commercially available viability/cytotoxicity assay kit (Biotium). After PBS washing, samples were exposed to staining medium containing 2 µM calcein AM and 4 µM EthD-III for 30 min at 37 °C. The stained samples were subsequently rinsed twice with PBS and immersed in fresh culture medium for fluorescence imaging. Viability was quantified from four independent image fields per sample collected at 20× magnification on a Leica DMi8 microscope.

### 4.9 Trans-epithelial electrical resistance measurement

TEER measurements were performed with a chopstick electrode (MERSSTX01, Millipore, USA) to assess the barrier properties of cell layers established on porous membranes. Net cellular resistance was determined by removing the contribution of the cell-free device from the measured resistance. The resulting value was multiplied by the membrane surface area to obtain TEER (Ω·cm²). Prior to TEER assessment of ALI cultures, 300 µL of medium was introduced into the apical chamber.

### 4.10 Immunofluorescence microscopy

For immunofluorescence analysis, cells cultured on the interstitium-mimicking membranes were rinsed three times with PBS and fixed in 4% paraformaldehyde (Sigma-Aldrich) for 20 min at room temperature. Following membrane sectioning, cells were permeabilized with 0.1% Triton X-100 for 10 min and subsequently blocked with 5% goat or donkey serum in PBS for 45 min. Samples were then incubated overnight at 4 °C with primary antibodies (Table S6) prepared in blocking solution. After washing, the corresponding secondary antibodies were applied for 1 h at room temperature. Nuclei were counterstained with DAPI, and fluorescence images were collected using a Zeiss LSM 880 confocal microscope (Germany).

### 4.11 Permeability measurement

Following removal of the culture medium, the device was rinsed with PBS. Phenol red-free medium (1 mL) was introduced into the basolateral compartment, while FITC–dextran (4 kDa, Sigma-Aldrich) solution (0.5 mL) was added to the apical compartment. Samples were collected from the basolateral outlet at 10 min intervals for 60 min while maintaining the device at 37 °C. Each collected sample was transferred to a black 96-well plate and analyzed only once. Fluorescence intensity was measured using a SpectraMax i3x plate reader with excitation and emission wavelengths of 490 and 525 nm, respectively.

The apparent permeability coefficient (P_app_, cm s-^1^) was determined using the equation below, where dQ/dt represents the molecular transport rate, A denotes the membrane area (cm^2^), and C_0_ corresponds to the initial FITC–dextran concentration in the apical compartment (mg mL⁻¹).

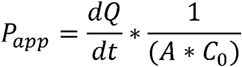

### 4.12 Quantification of apoptosis, cell density, and **γ**-H2AX positivity

To assess cell density, DAPI-positive nuclei were quantified within a defined area and normalized to the corresponding surface area. Apoptosis was evaluated by determining the fraction of cleaved caspase-3-positive cells among the total cell population. The percentage of γ-H2AX-positive cells was calculated in an analogous manner. Data were obtained from four randomly selected images per sample and reported as the mean value.

### 4.13 Measurement of Hydrogen Peroxide, 8-OHdG, and Cytokines

Hydrogen peroxide (HLJOLJ), 8-OHdG, and cytokine levels were quantified using a Hydrogen Peroxide Assay Kit (Sigma-Aldrich), a DNA Damage ELISA Kit (StressMarq Biosciences), and a Multiplex Human Cytokine ELISA Kit (Anogen), respectively, following the suppliers’ instructions. To account for background contributions, normalized HLJOLJ production was calculated as H_A_ − H_B_ − H_C_, where HA corresponds to the signal obtained from coal dust-exposed cultures treated with vitamins, H_B_ denotes the baseline HLJOLJ level produced by cells cultured on the chip in the absence of coal dust and vitamins, and H_C_ represents the signal attributable to vitamin treatment alone.

### 4.14 Polycyclic aromatic hydrocarbons (PAHs) measurement from coal sample

PAHs were extracted from the coal sample following U.S. EPA Method 3540C (Soxhlet Extraction) [92]. Prior to extraction, coal samples were air-dried, homogenized, and sieved to obtain a representative particle size fraction. Representative coal sample was initially cryo-milled for 2 cycles at 30 Hz, with each 2 min milling each cycle. This procedure produced finely divided, coal particles suitable for subsequent chemical extraction and characterization analyses. Approximately 5 g prepared sample was mixed with anhydrous sodium sulfate to remove residual moisture and transferred into cellulose extraction thimbles. Soxhlet extraction was performed using an organic solvent system consistent with EPA 3540C, typically dichloromethane or a dichloromethane–acetone mixture, to ensure efficient recovery of both low- and high-molecular-weight PAHs. Samples were extracted for a minimum of 16 hours under continuous reflux conditions to achieve exhaustive extraction of semi-volatile organic compounds. Following extraction, the solvent extracts were allowed to cool and were subsequently concentrated using rotary evaporation and gentle nitrogen blowdown to a final volume suitable for instrumental analysis. Extracts were subjected to cleanup procedures, as required, to remove co-extracted matrix interferences prior to analysis. Cleanup steps included silica or alumina column chromatography in accordance with EPA guidelines. Final extracts were stored in amber glass vials at 4 °C until chemical analysis.

Identification and quantification of PAHs were conducted using U.S. EPA Method 8270E, employing gas chromatography–mass spectrometry (GC–MS) operated in electron ionization (EI) mode [93]. Analyses were performed using a capillary GC column suitable for semi-volatile organic compounds. Instrument calibration was achieved using multi-point calibration curves generated from certified PAH standard mixtures spanning the expected concentration range of the samples. Target analytes were identified based on retention time matching and characteristic mass spectral fragmentation patterns compared to authentic standards and reference libraries. Quantification was performed using internal standard calibration, with surrogate standards added prior to extraction to evaluate method recovery and analytical performance. Method blanks, matrix spikes, and duplicate samples were analyzed periodically to ensure data quality and compliance with QA/QC requirements specified in EPA Method 8270E. Detection limits and quantification limits were determined according to EPA guidelines, and all reported concentrations were blank-corrected where applicable. PAH concentrations are reported on a dry-weight basis.

### 4.15 Bulk RNA sequencing

To remove residual coal dust, the cells were washed five times with PBS. Cells from each chip were harvested by trypsinization, and two chips were pooled to obtain a sufficient amount of RNA for each sample. RNA extraction, library preparation and sequencing were performed by Novogene Corporation, Inc. Briefly, total RNA was extracted and used for directional mRNA library preparation employing poly(A) enrichment. Libraries were sequenced on an Illumina NovaSeq X Plus platform to generate 150 bp paired-end reads, yielding approximately 9 Gb of raw data per sample. Raw sequencing reads were evaluated for quality using FastQC (v0.11.9). Adapter sequences and low-quality bases were removed using Trim Galore (v0.6.10) with default parameters. Trimmed reads were quantified using Salmon (v1.10.3) in quasi-mapping mode with GC bias correction enabled [94]. A Salmon index was built from the human reference transcriptome (GENCODE v49, based on GRCh38.p14) using the entire GRCh38 genome as a decoy sequence. Mapping files output by Salmon were further quality-controlled using Picard (v2.25.1) to assess alignment metrics. Transcript-level quantifications were summarized to gene-level counts using the tximport (v1.32.0) package in R and analyzed using DESeq2 (v1.46.0) [95]. Genes with fewer than ten total counts in at least three samples were excluded prior to normalization. Size factors and dispersion estimates were computed using DESeq2’s standard workflow. Differential expression (DE) between experimental groups was tested using the Wald test, and p-values were adjusted for multiple testing using the Benjamini–Hochberg false discovery rate (FDR) procedure. logD fold changes were shrunk using the adaptive shrinkage estimator ash [96]. Differential expression results were visualized using the EnhancedVolcano (v1.24.0) R package. Gene set enrichment analysis (GSEA) was performed using the fgsea (v1.32.2) R package. Genes were ranked by signed p-values derived from the differential expression analysis, calculated as sign(log2(fold change)) × –log10(p-value). Gene sets were obtained from the MSigDB database using the msigdbr (v25.1.1) R package. P-values were adjusted for multiple testing using the Benjamini–Hochberg false discovery rate (FDR) procedure. Heatmaps of leading-edge genes were generated using the pheatmap (v1.0.13) R package.

### 4.16 Statistics

Student’s t-test was employed to evaluate differences between two groups. When three or more groups were compared, one-way or two-way ANOVA with Tukey’s post hoc correction was used. A p value of less than 0.05 was considered indicative of statistical significance.

## Acknowledgements

This study was financially supported by US National Institute for Occupational Safety and Health (NIOSH, 75D30119C05128), and the Philip and Marsha Dowd Fellowship of Carnegie Mellon University.

## Data Availability Statement

The data that support the findings of this study are available from the corresponding author upon reasonable request.

