## Supplemental File for "Interstitium-mimicking porous alveolar membranes enable physiologic aerosol transport and distinct acute-chronic lung injury responses"

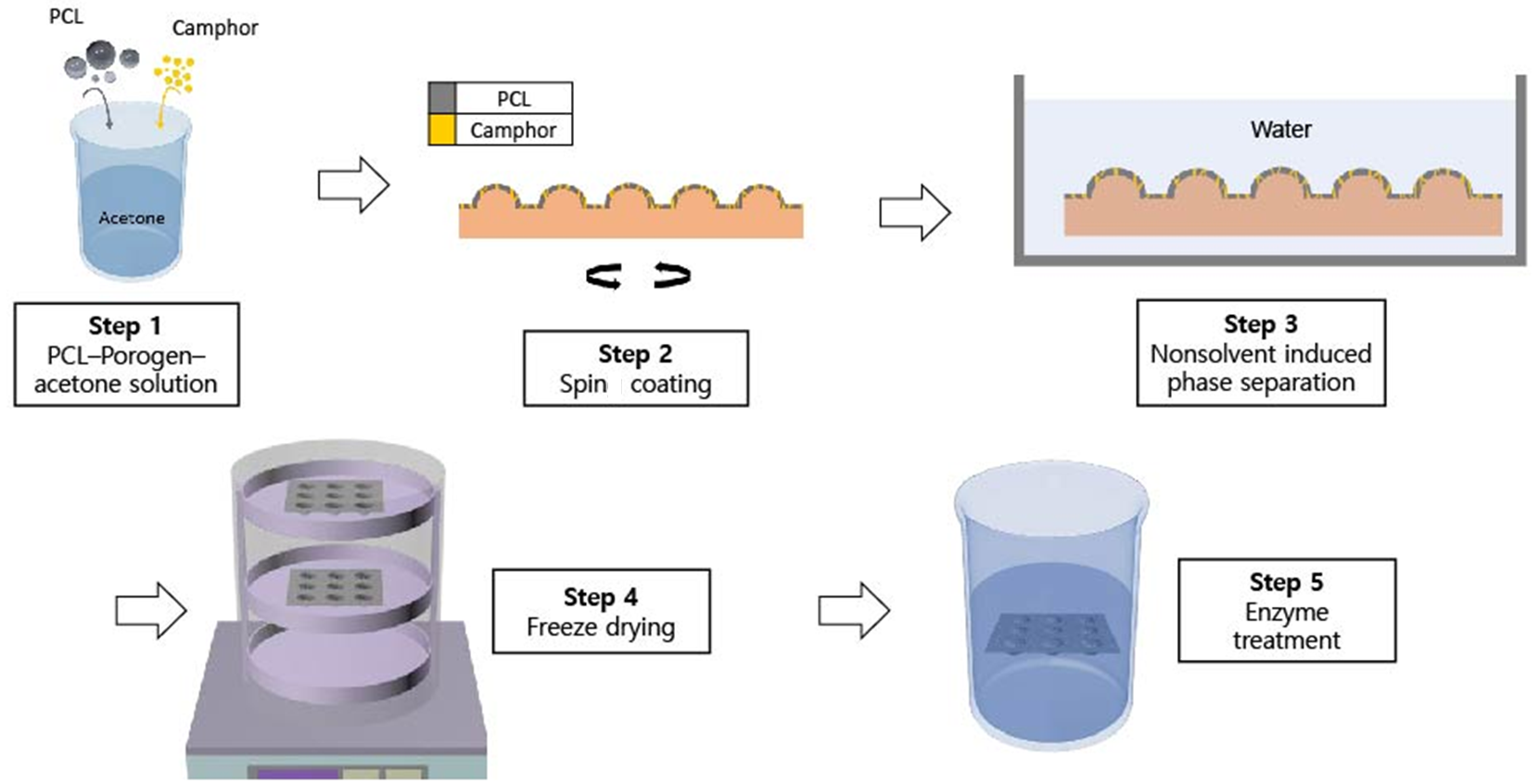

**Figure S1.** Schematic illustration of the fabrication process of the alveolus-shaped porous PCL membrane using dual-templated nonsolvent-induced phase separation and subsequent lipase-mediated pore enlargement.

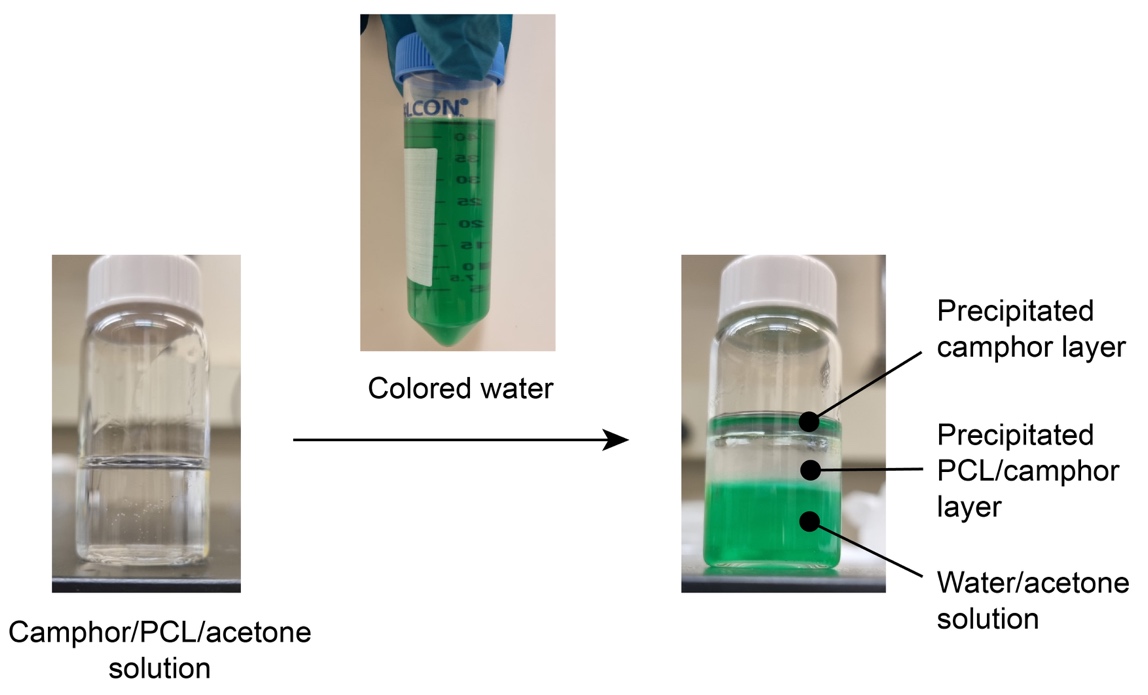

**Figure S2.** Optical images of the camphor–PCL solution in acetone (left) and phase separation upon immersion in water/acetone mixture (right), showing precipitation of camphor and formation of the PCL/camphor composite layer.

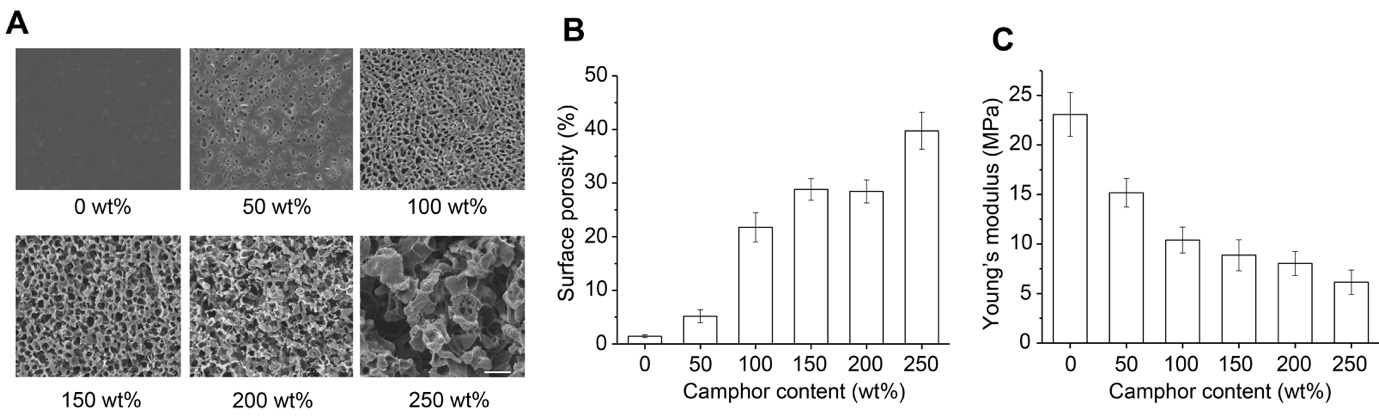

**Figure S3.** Effect of camphor loading on pore architecture and mechanical properties of porous PCL membranes. (A) SEM images showing surface pore morphology as a function of camphor content (0, 50, 100, 150, 200, and 250 wt% relative to PCL). Scale bar: 20 µm. (B) Surface porosity and (C) Young’s modulus of porous PCL membranes as a function of camphor content (n = 4).

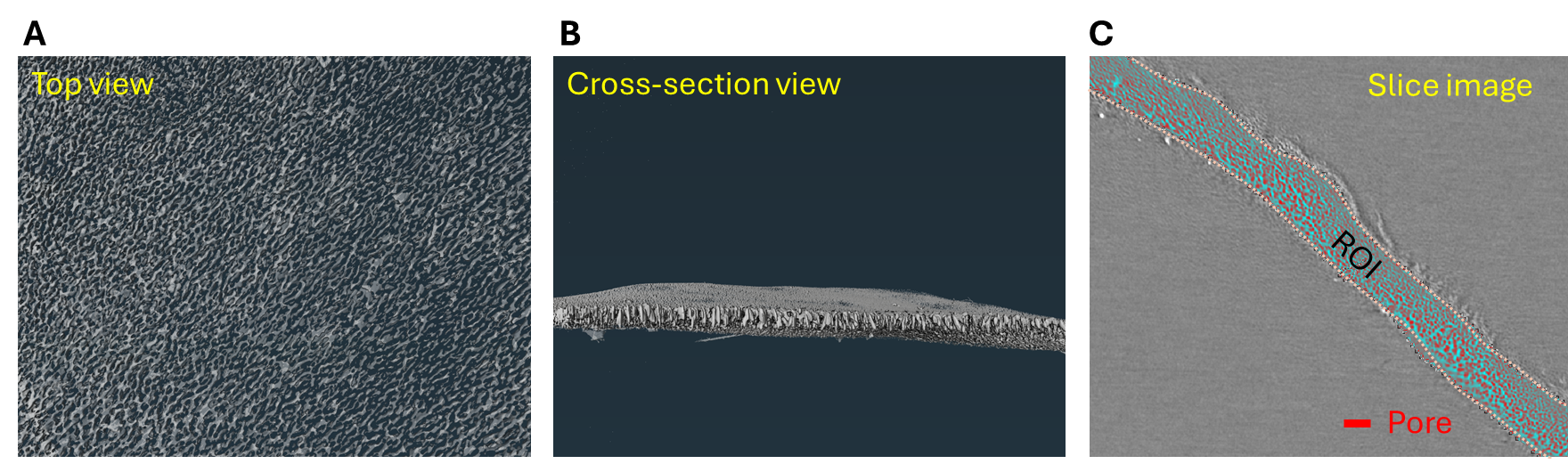

**Figure S4.** Representative nano-CT analysis of the porous PCL membrane. (A) Top-view 3D reconstruction showing the surface porous morphology. (B) 3D cross-sectional reconstruction showing the through-thickness architecture of the membrane. (C) A reconstructed slice image showing the manually defined region of interest (ROI) and segmented pore space.

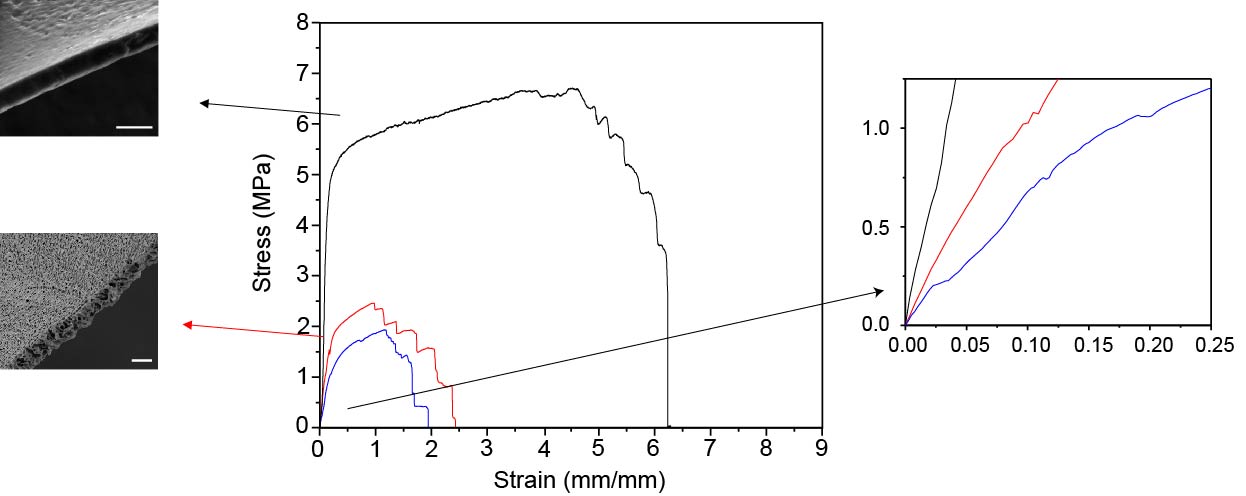

**Figure S5.** Representative stress-strain curves of nonporous PCL membrane and alveolus-shaped porous PCL membranes before and after lipase treatment during tensile testing (black: nonporous; red: porous before lipase treatment; blue: porous after lipase treatment).

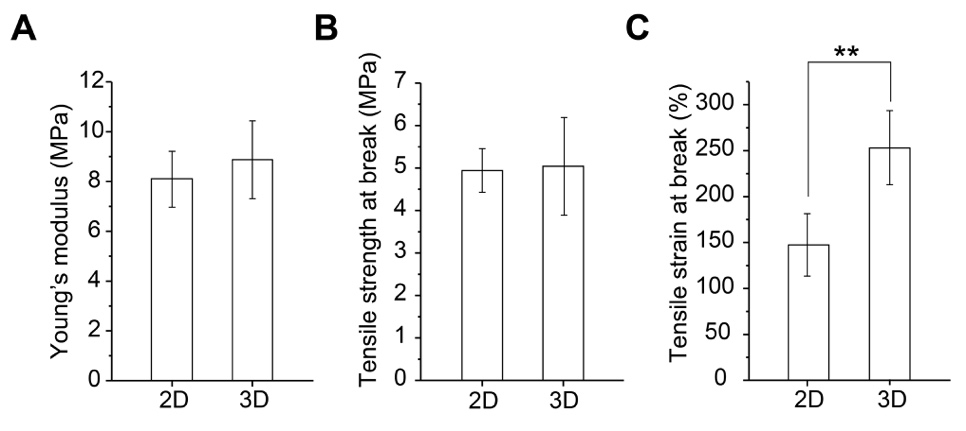

**Figure S6.** Comparison of mechanical properties between flat (2D) porous PCL membranes and alveolus-shaped (3D) porous PCL membranes prior to lipase treatment: (A) Young’s modulus, (B) tensile strength, and (C) tensile strain at break (n = 5).

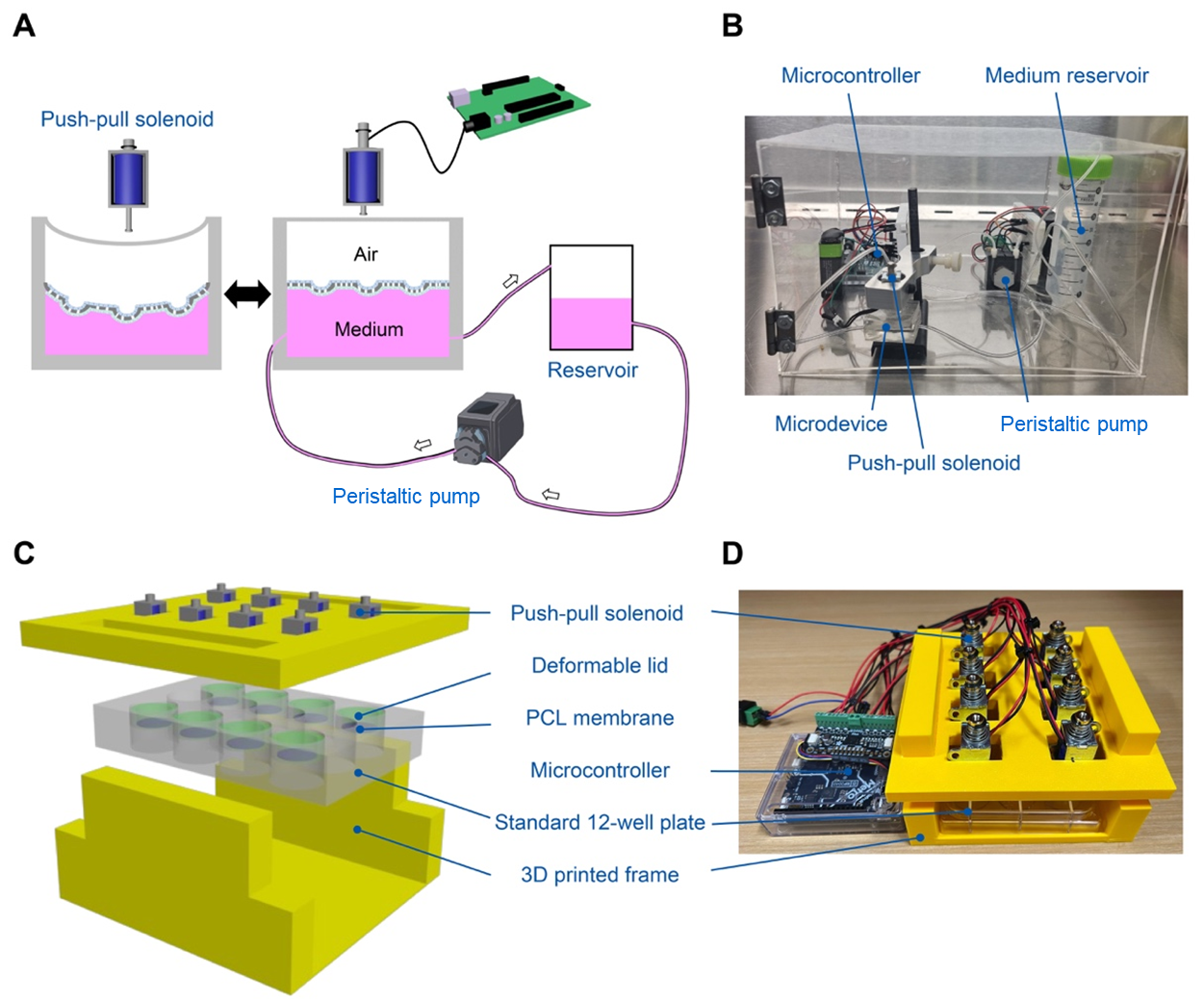

**Figure S7.** Configuration of the alveoli-on-chip system. (A) Schematic of a single device showing the compartmentalized architecture. The basolateral chamber is connected to inlet and outlet reservoirs for continuous perfusion via a peristaltic pump, while a microcontroller-driven actuator applies cyclic deformation to the apical lid, transmitting strain to the interstitium-mimicking PCL membrane. (B) Photograph of a single alveoli-on-chip device. (C) Schematic of an eight-unit array configured in a standard 12-well plate format. (D) Photograph of the eight-unit array.

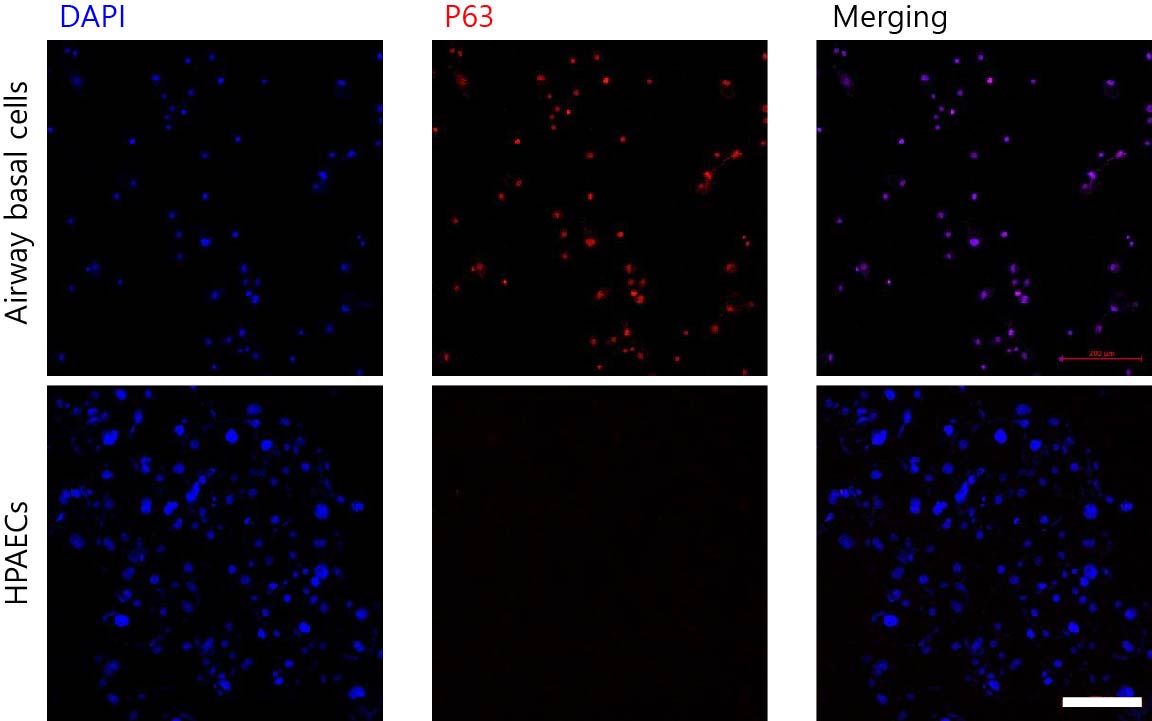

**Figure S8.** Immunofluorescence staining for P63, a basal cell marker. Airway basal stem cells (upper panel) show positive P63 expression, whereas primary human alveolar epithelial cells (HPAECs, lower panel) show negligible expression, confirming alveolar phenotype. Cells were cultured in 24-well plates and fixed 48 h after seeding. Scale bars: 200 µm.

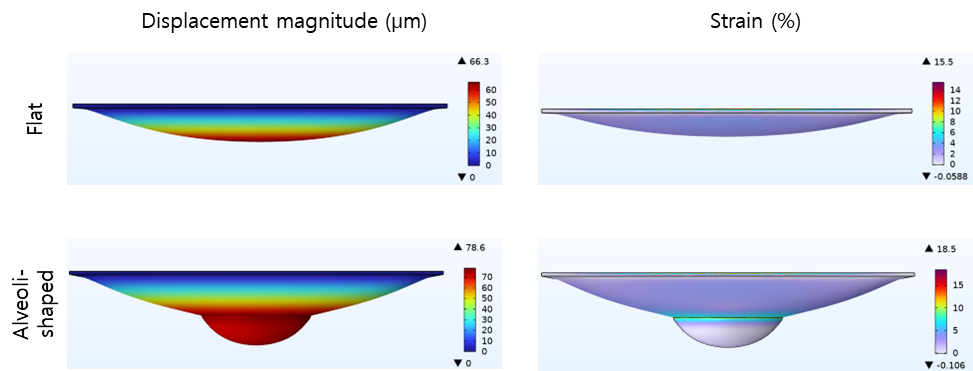

**Figure S9.** Finite element simulation comparing deformation of flat and alveolus-shaped membranes under identical applied pressure (20 mbar). The top row shows the flat membrane, and the bottom row shows the alveolus-shaped membrane. Displacement magnitude (right) and strain distribution (left) are displayed for each geometry.

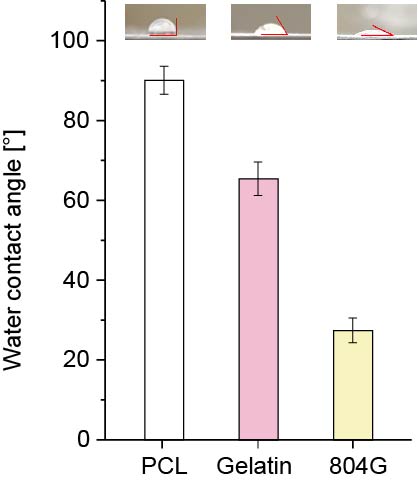

**Figure S10.** Water contact angle measurements of bare porous PCL membranes and membranes coated with gelatin or 804G-conditioned medium (n = 4).

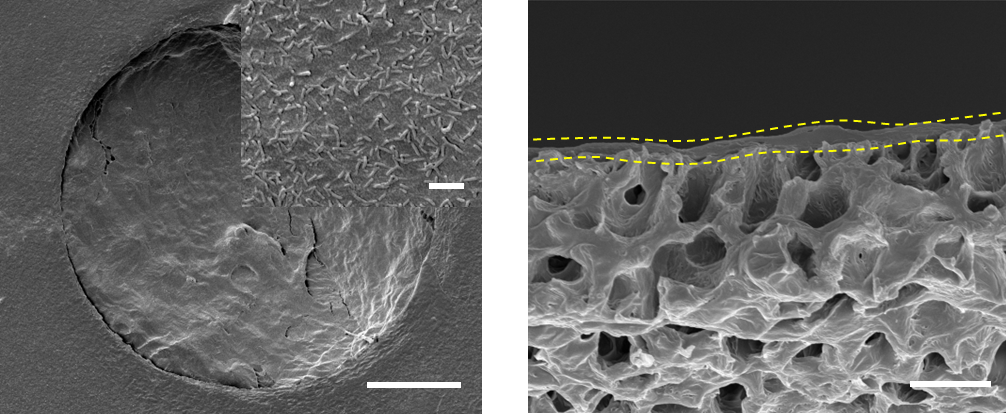

**Figure S11.** SEM characterization of airway basal stem cells cultured on the alveolus-shaped porous PCL membrane. Left: Representative top-view SEM image showing airway basal stem cells on the apical surface (scale bar: 50 µm), with inset highlighting ciliated cells after 21 days of ALI differentiation (scale bar: 1 µm). Right: Cross-sectional SEM image showing a confluent monolayer of airway basal stem cells on the apical side of the membrane (scale bar: 5 µm).

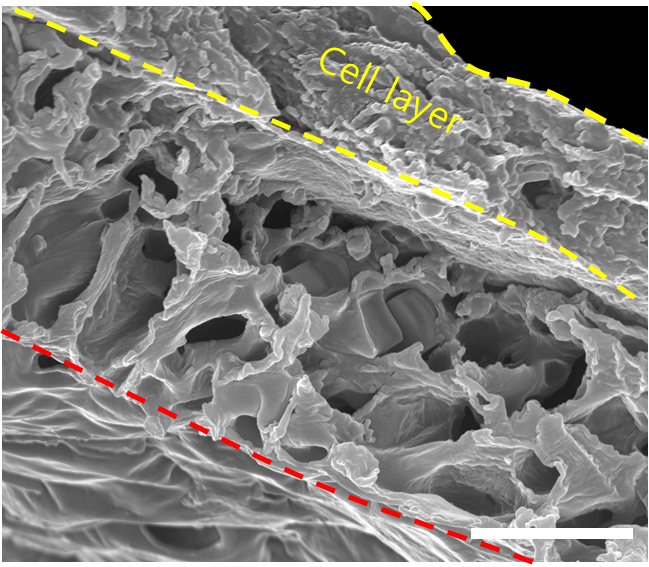

**Figure S12.** Representative cross-sectional SEM image of A549 cells cultured on the alveolus-shaped porous PCL membrane at day 8 (7 days in LLC followed by 24 h in ALI). A confluent epithelial monolayer is formed on the apical surface, with media supplied from the basolateral chamber through the vertically interconnected pore network. Scale bar: 5 µm.

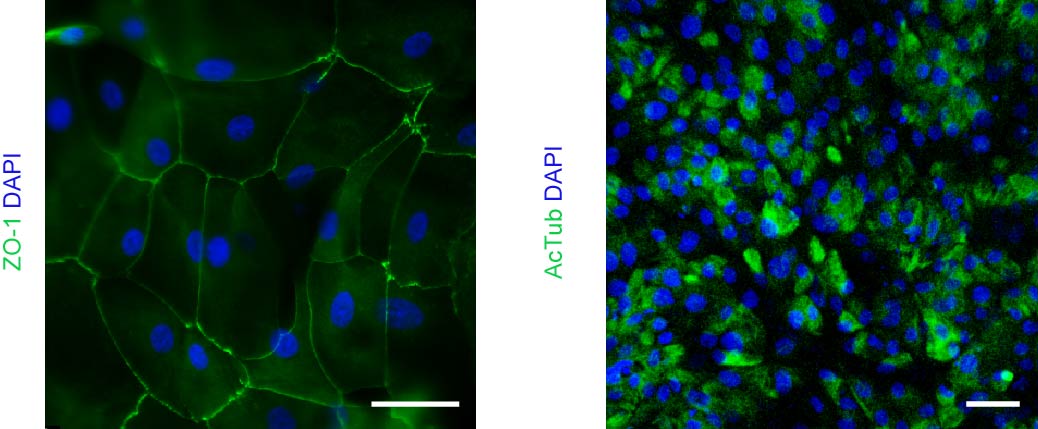

**Figure S13.** Confocal fluorescence images of airway basal stem cell cultures on the alveolus-shaped membrane. Left: Confluent epithelial monolayer with tight junction protein ZO-1 (green) and nuclei (DAPI, blue). Right: Differentiated ciliated cells stained for acetylated tubulin (green) with nuclei (DAPI, blue). Scale bars: 50 µm.

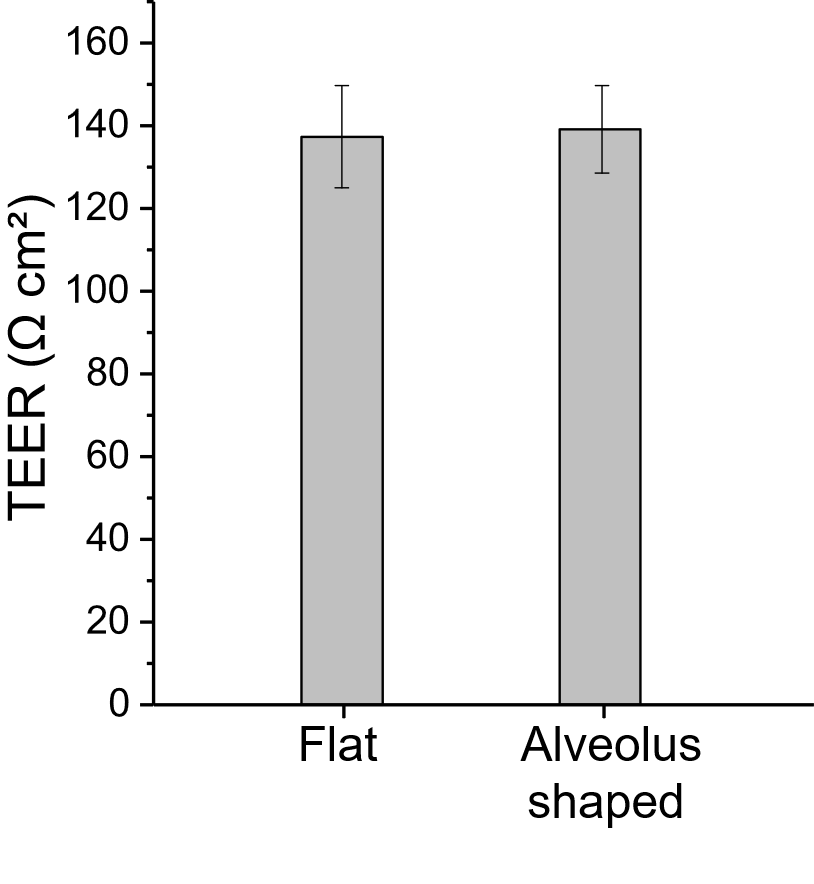

**Figure S14.** TEER of A549 cell monolayers cultured on flat and alveolus-shaped membranes. The alveolus-shaped membrane consisted of both flat (2D) and dome-shaped (3D) regions with an approximate surface area ratio of 7:1. TEER values (Ω·cm²) were calculated by multiplying the measured cell resistance (Ω) by the total membrane surface area, including the 3D dome-shaped regions (n = 4).

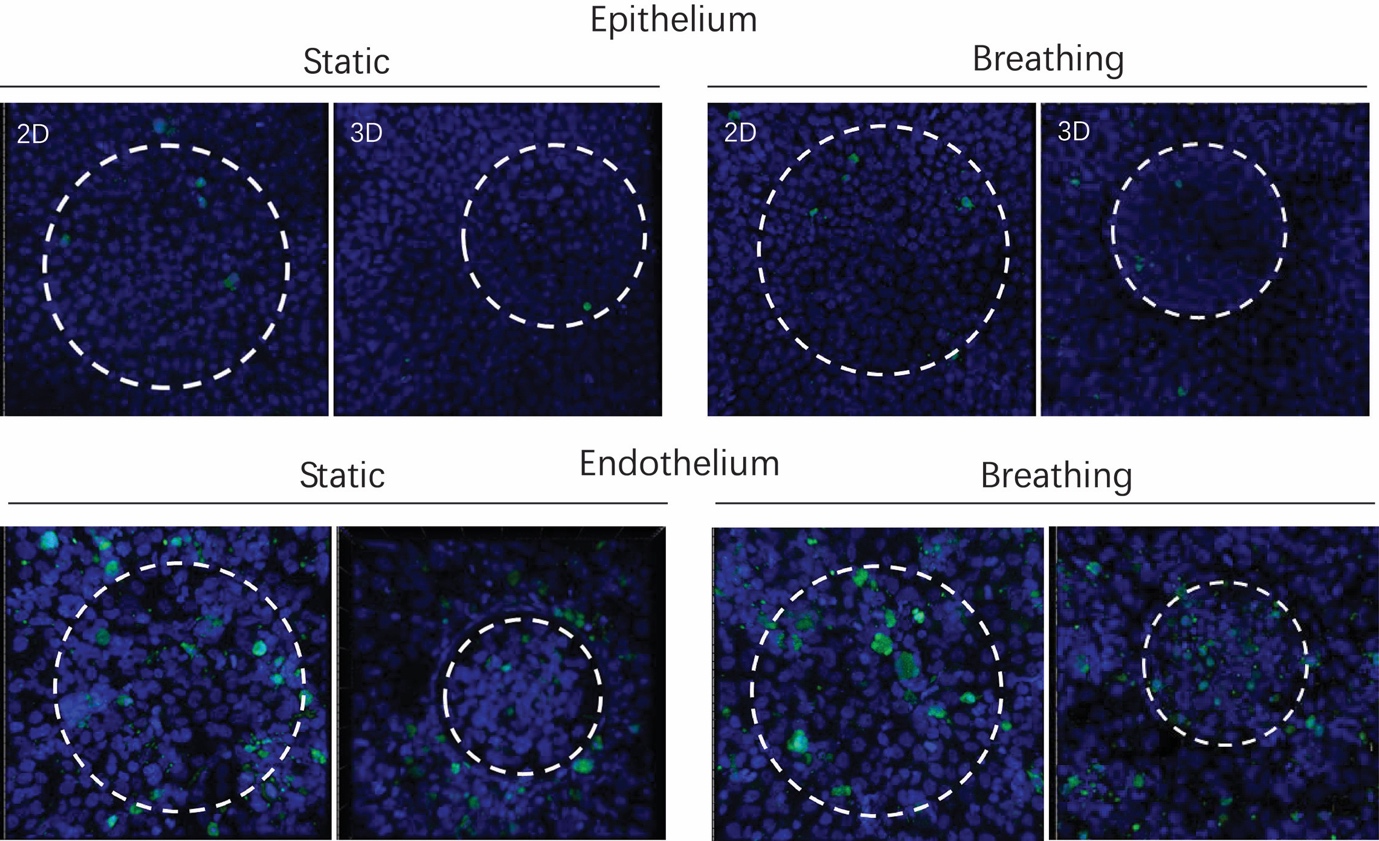

**Figure S15.** Representative fluorescence images of apoptosis in A549 cells (top) and HUVECs (bottom) under static and cyclic deformation conditions. Cells were stained for cleaved caspase-3 (green) and nuclei (DAPI, blue). White dashed circles indicate equivalent surface area between flat (2D) and hemispherical alveolus-shaped (3D) regions used for comparison. Scale bars: 100 µm.

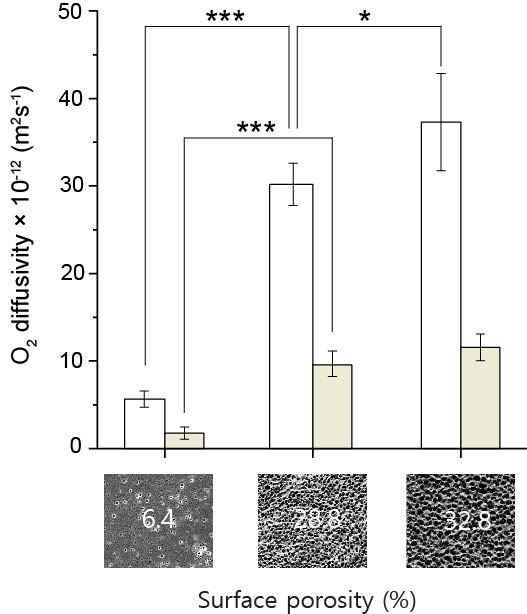

**Figure S16.** Oxygen diffusivity of porous PCL membranes with varying porosity (6.4%, 28.8% before lipase treatment, and 32.8% after lipase treatment) under acellular and epithelial–endothelial co-culture conditions. In cellular conditions, A549 cells and HUVECs were cultured on opposite sides of the membrane. White bars: acellular; beige bars: cellular (n = 4).

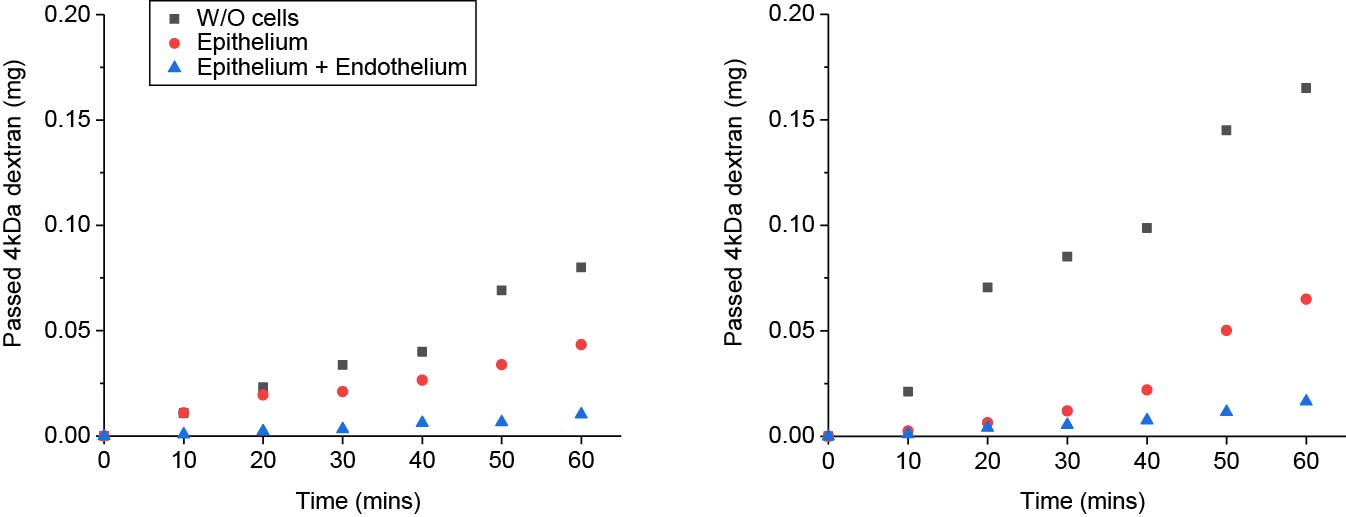

**Figure S17.** Time-dependent transport of 4 kDa FITC-dextran across the alveolus-shaped porous PCL membrane and A549-HUVEC co-culture layers under static (left) and cyclic deformation (right) conditions.

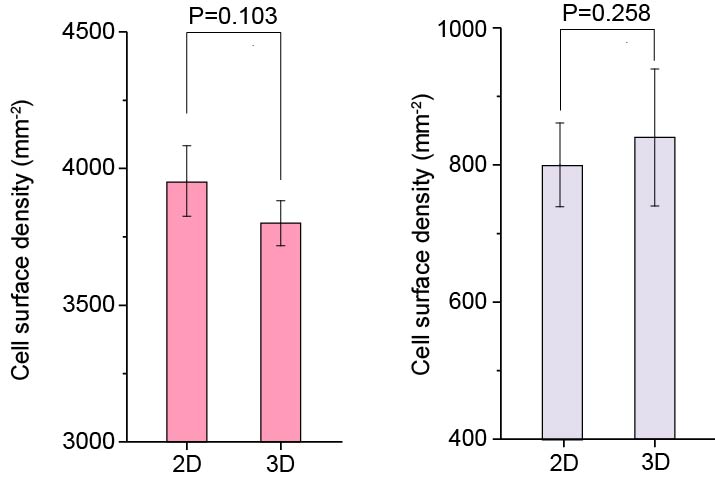

**Figure S18.** Surface cell density of HPAECs (left) and HMVECs (right) under cyclic deformation. Cell densities were compared between flat (2D) regions and hemispherical alveolus-shaped (3D) regions (n = 5).

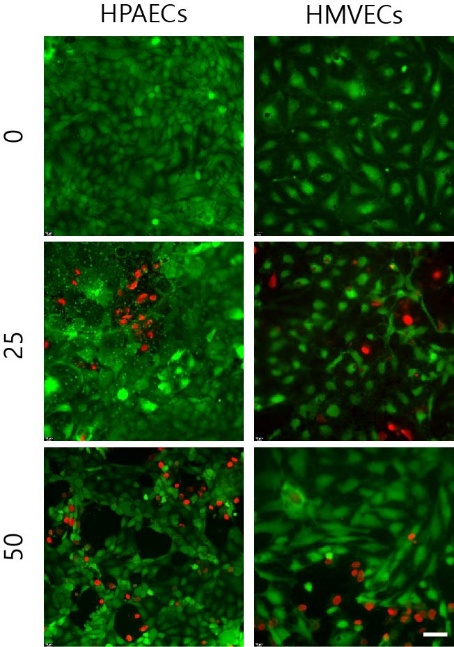

**Figure S19.** Representative images of HPAEC-HMVEC co-cultures in the alveoli-on-chip system 24 h after coal dust exposure at surface loadings of 0, 25, and 50 µg cm⁻². Scale bars: 50 µm.

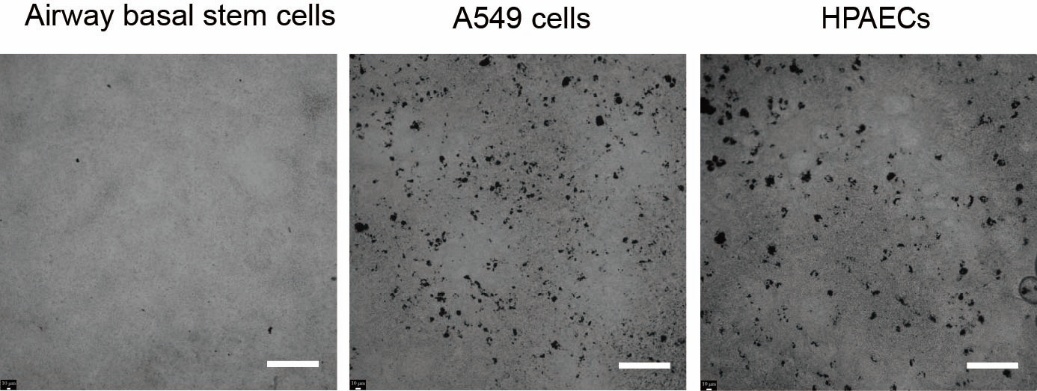

**Figure S20.** Representative bright-field images showing coal dust adherence to airway basal stem cells (left), A549 cells (middle), and HPAECs (right) 24 hours after exposure. Cells were cultured on flat porous PCL membranes prior to particulate exposure. Scale bars: 100 µm.

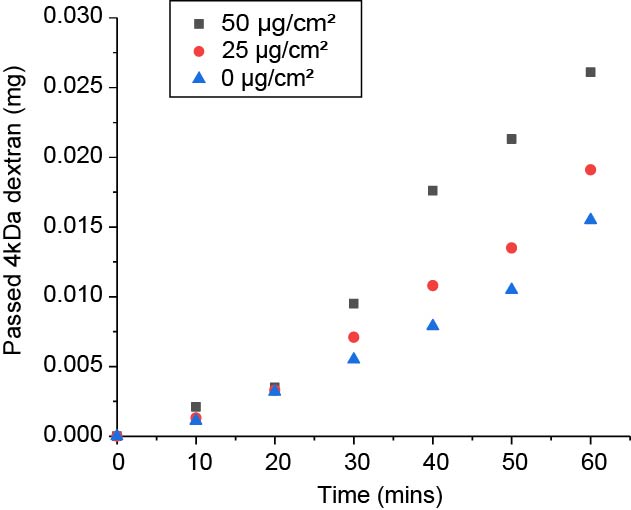

**Figure S21.** Time-dependent permeability of 4 kDa FITC-dextran across A549-HUVEC co-cultures measured 24 hours after coal dust exposure at surface loadings of 0, 25, and 50 µg cm⁻².

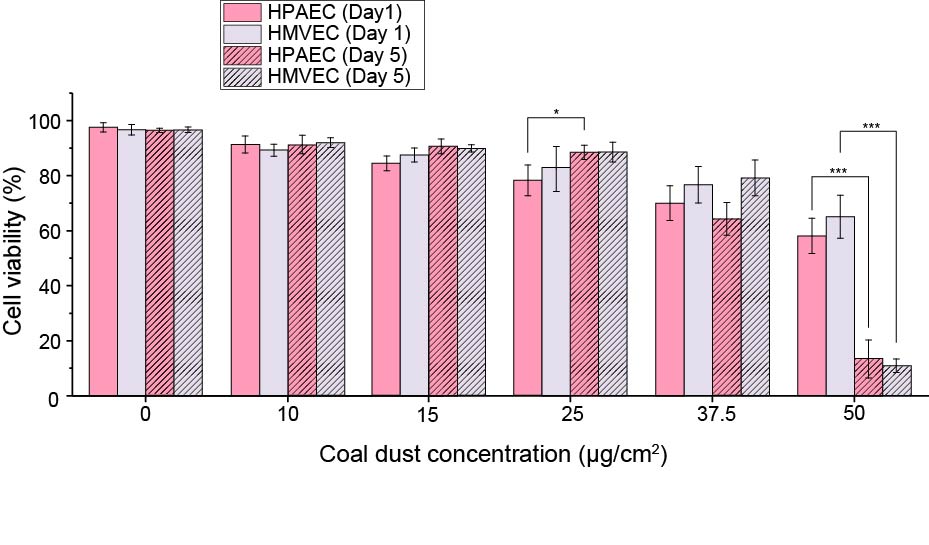

**Figure S22.** Dose-dependent effects of coal dust on HPAEC–HMVEC co-culture viability over time. Cell viability was measured on days 1 and 5 following exposure to surface loadings of 0, 10, 15, 25, 37.5, and 50 µg cm⁻² (n = 4).

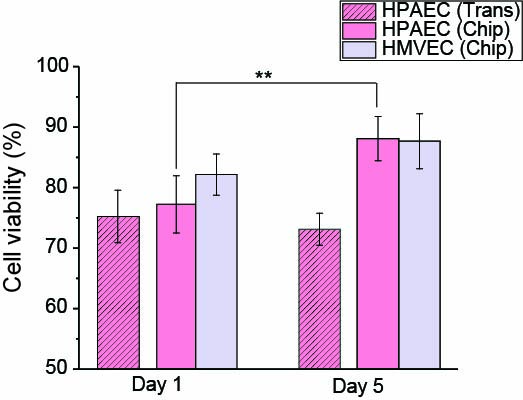

**Figure S23.** Comparison of epithelial and endothelial cell viability after coal dust exposure in Transwell® cultures (HPAEC monoculture) and alveoli-on-chip co-cultures (HPAEC-HMVEC). ALI culture was initiated on day 7; cyclic deformation (10% linear strain, 0.33 Hz) was applied in the chip system from day 7 onward. Coal dust (25 µg cm⁻²) was applied on day 8, and cell viability was assessed on days 1 and 5 post-exposure (n = 4).

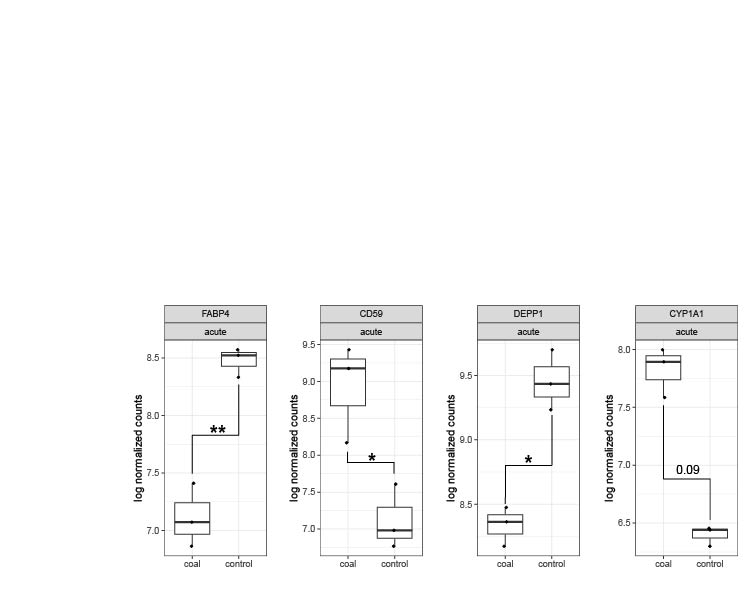

**Figure S24.** Differential expression of selected genes in coal dust-treated versus untreated control samples under acute exposure conditions (n = 3).

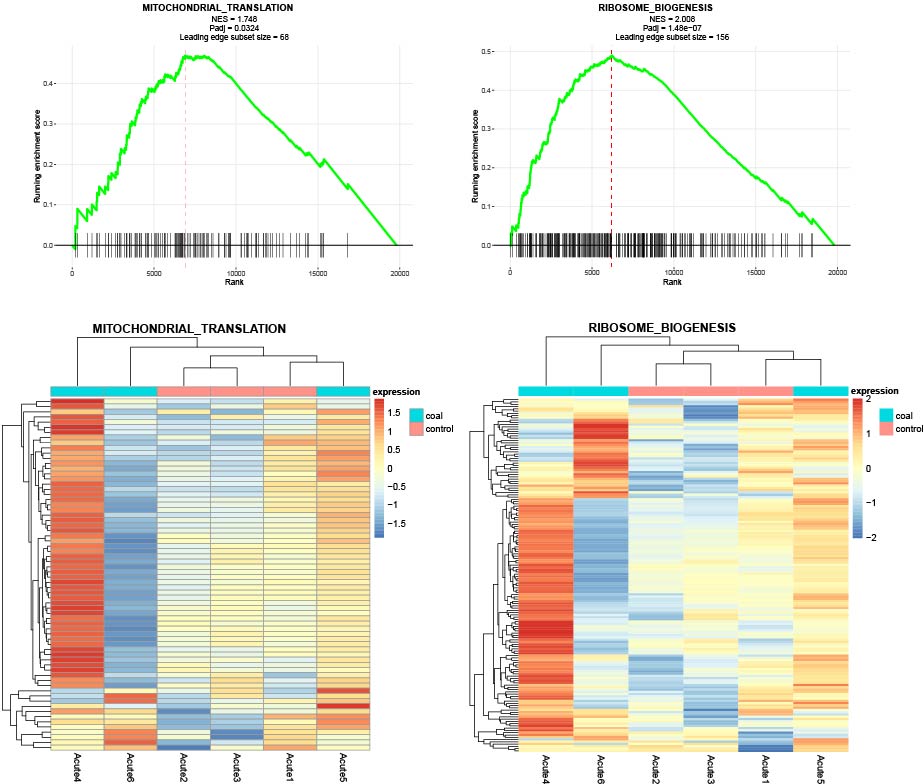

**Figure S25.** GSEA results and leading-edge gene heatmaps for additional significantly altered pathways under acute coal dust exposure.

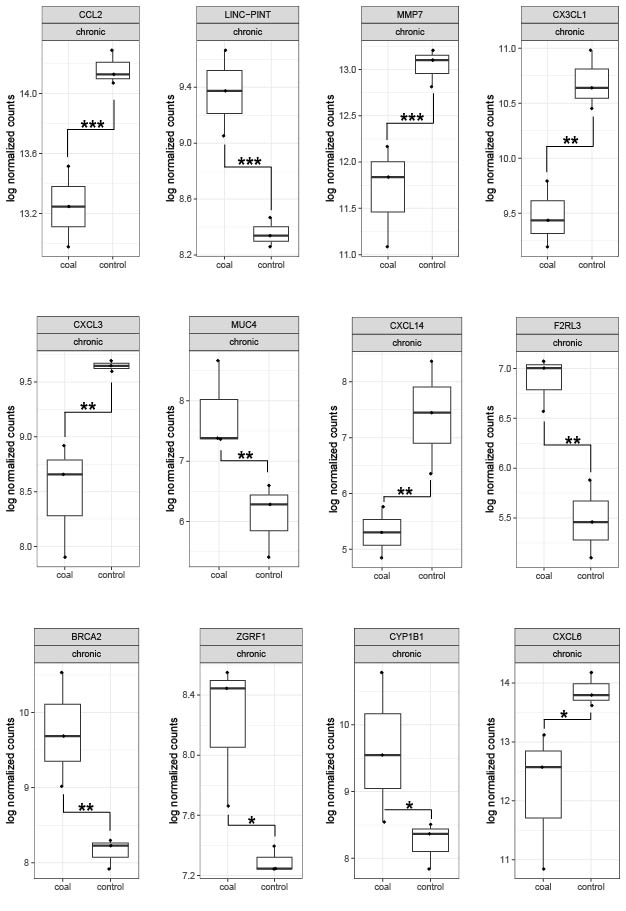

**Figure S26.** Differential expression of selected genes in coal dust-treated versus untreated control samples under chronic exposure conditions (n = 3).

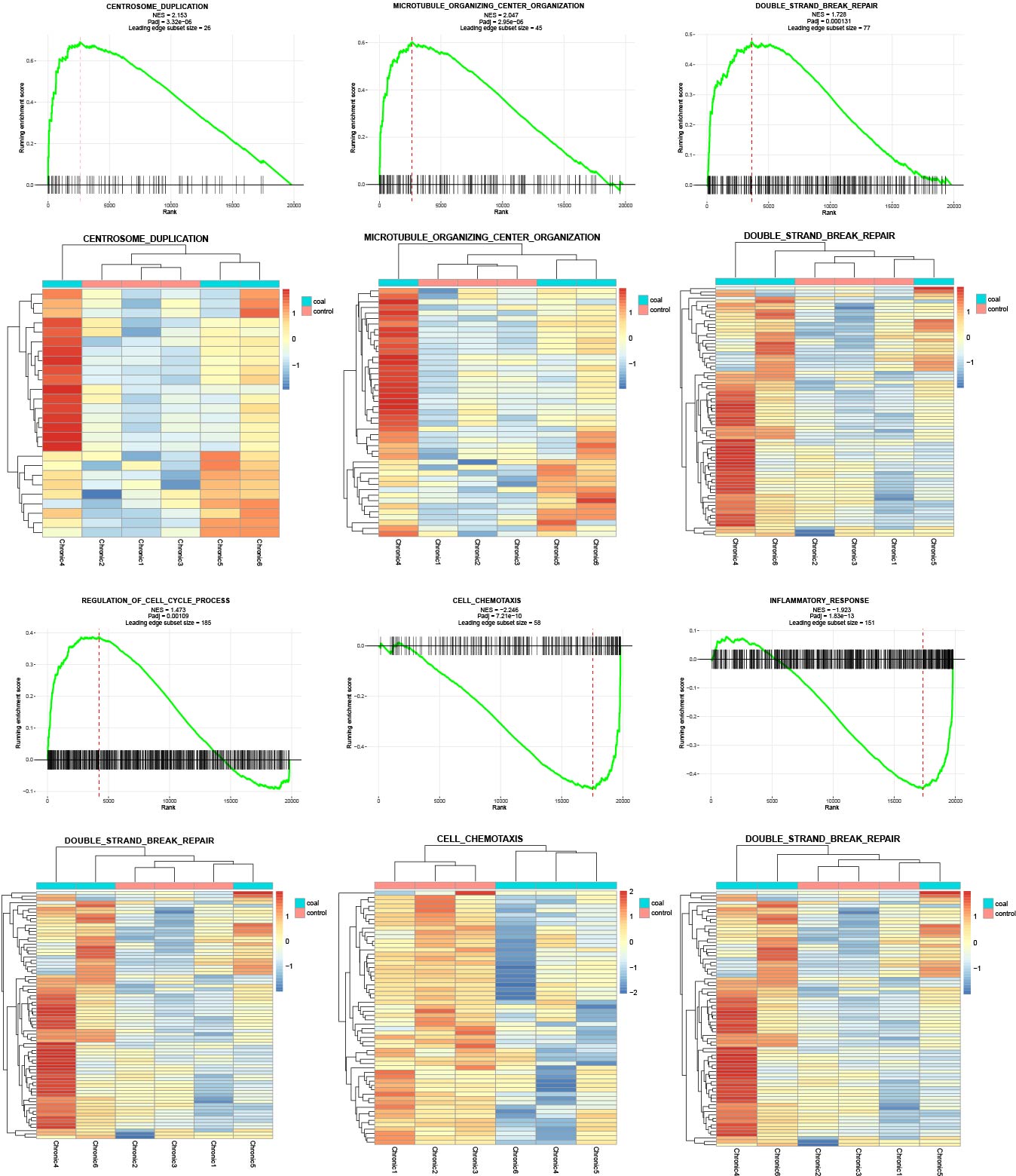
 **Figure S27.** GSEA results and leading-edge gene heatmaps for additional significantly altered pathways under chronic coal dust exposure.

|  | Material | Dimension | Porosity  (%) | Thickness  (μm) | Pore size (µm) | Cyclic pressure | Epithelium | Endothelium | ALI culture | Application |
| --- | --- | --- | --- | --- | --- | --- | --- | --- | --- | --- |
| CMU (this work) | PCL | 3D | 40 | 2.5 | 3~4 | 10%, 0.33Hz | ✓ | ✓ | ✓ | Nano coal dust |
| Advanced functional materials (2021) [1] | PCL | 2D | 9.4 | 5 | 4.5 | 10%, 0.33Hz | ✓ | ⎯ | ✓ | Nano, Microparticles |
| Advanced materials (2022) [2] |  |  | 20 | 1 |  | 10%, 0.33Hz | ✓ | ✓ | ✓ | Lung fibrosis |
| Nature BME (2021) [3] | PET | 2D | 8.8 | 10 | 0.4 | ⎯ | ✓ | ✓ | ✓ | Influenza virus |
| Nature Communications (2022) [4] | PDMS  (Emulate Inc) | 2D | 2.5 | 50 | 7 | 5%, 0.25Hz | ✓ | ✓ | ✓ | Influenza virus |
| Nature Communications (2023) [5] |  |  |  |  |  |  |  |  |  | Acute radiation-induced lung injury |
| PNAS (2021) [6] | Gelatin methacryloyl | 3D | ⎯ | ⎯ | ⎯ | 10%, 0.2Hz | ✓ | ⎯ | ✓ | Smoking |
| Biomaterials (2021) [7] | Polycarbonate | 3D | <5 | 10 | 4.6 | ⎯ | ✓ | ⎯ | ✓ | ⎯ |
| Nature BME (2025) [8] | PET | 2D | 0.25 |  | 0.4 | ⎯ | ✓ | ✓ | ✓ | SARS-CoV-2  Influenza virus |
| Nature BME (2025) [9] | PET | 2D | 7.9 | 10 | 5 | ⎯ | ✓ | ✓ | ✓ | Influenza virus |

**Table S1.** Summary of membrane materials and properties used in representative lung-on-chip platforms. Abbreviations: PCL, poly(ε-caprolactone); PET, polyethylene terephthalate; PDMS, polydimethylsiloxane.

| Camphor ratio (wt%) | 0 | 50 | 100 | 150 | 200 | 250 |
| --- | --- | --- | --- | --- | --- | --- |
| Porosity (%) | 1.45 | 5.17 | 21.74 | 28.81 | 28.45 | 39.74 |
| Young’s modulus (MPa) | 23.09 | 15.17 | 10.41 | 8.87 | 8.04 | 6.14 |

**Table S2.** Effect of camphor loading (wt% relative to PCL) on membrane porosity and Young’s modulus.

|  | Nonporous structure | Alveoli-shaped  (Before enzyme treatment) | Alveoli-shaped  (After enzyme treatment) |
| --- | --- | --- | --- |
| Young’s modulus (MPa) | 23.09 ± 2.22 | 8.87 ± 1.57 | 5.34 ± 1.17 |
| Yield strength (MPa) | 5.68 ± 0.23 | 1.95 ± 0.42 | 1.74 ± 0.20 |
| Ultimate tensile strength (MPa) | 6.62 ± 0.27 | 2.52 ± 0.57 | 1.97 ± 0.26 |

**Table S3.** Comparison of mechanical properties (Young’s modulus, yield strength, and ultimate tensile strength) of nonporous PCL membranes and alveolus-shaped porous PCL membranes before and after lipase treatment.

|  | PLC | Transwell® | PCL | PCL |
| --- | --- | --- | --- | --- |
| Porosity (%) | 6.4 | 9.1 | 28.8 | 32.8 |
| O_2_ diffusivity × 10^-11^  (m^2^s^-1^) | 0.564 ± 0.092 | 0.698 ±  0.130 | 3.018 ± 0.242 | 3.729 ±  0.555 |

**Table S4.** Quantification of oxygen diffusivity across interstitium-mimicking membranes with surface porosities of 6.4%, 28.8%, and 32.8%, and a Transwell® membrane with a surface porosity of 9.1% (n = 4).

| Pathway | padj | NES | Leading Edge |
| --- | --- | --- | --- |
| Acute Condition | | | |
| RIBOSOME BIOGENESIS | 0.000000147889 | 2.0079 | 156 |
| RIBONUCLEOPROTEIN COMPLEX BIOGENESIS | 0.000000259344 | 1.8831 | 221 |
| TELOMERE MAINTENANCE VIA TELOMERE LENGTHENING | 0.044228752017 | 1.8001 | 71 |
| MITOCHONDRIAL TRANSLATION | 0.032351583759 | 1.7477 | 68 |
| RNA PROCESSING | 0.000000204724 | 1.6460 | 389 |
| TRNA METABOLIC PROCESS | 0.036683590432 | 1.5883 | 78 |
| CHROMOSOME ORGANIZATION | 0.033005971003 | 1.4352 | 166 |
| DNA METABOLIC PROCESS | 0.036683590432 | 1.3129 | 245 |
| CELL-CELL ADHESION | 0.000010758098 | -1.6362 | 210 |
| INFLAMMATORY RESPONSE | 0.000013097573 | -1.6950 | 182 |
| LOCOMOTION | 0.000000041811 | -1.7037 | 253 |
| VASCULATURE DEVELOPMENT | 0.000000147889 | -1.7874 | 202 |
| LEUKOCYTE MIGRATION | 0.000137459033 | -1.8528 | 60 |
| TAXIS | 0.000030808784 | -1.8665 | 105 |
| REGULATION OF VASCULAR PERMEABILITY | 0.033588315798 | -1.8693 | 15 |
| CELL-CELL ADHESION VIA PLASMA MEMBRANE ADHESION MOLECULE | 0.000026630153 | -2.0220 | 61 |
| Chronic Condition | | | |
| CENTROSOME DUPLICATION | 0.000003323892 | 2.1535 | 26 |
| MICROTUBULE ORGANIZING CENTER ORGANIZATION | 0.000002950113 | 2.0475 | 45 |
| DNA TEMPLATED DNA REPLICATION MAINTENANCE OF FIDELITY | 0.003813368211 | 1.9488 | 25 |
| ORGANELLE FISSION | 0.000000240354 | 1.8441 | 101 |
| CHROMOSOME ORGANIZATION | 0.000000006709 | 1.8363 | 120 |
| DOUBLE STRAND BREAK_REPAIR | 0.000130895186 | 1.7282 | 77 |
| MICROTUBULE CYTOSKELETON ORGANIZATION | 0.000000240354 | 1.7158 | 141 |
| DNA REPLICATION | 0.002179401248 | 1.6476 | 106 |
| DNA REPAIR | 0.000054269735 | 1.5813 | 204 |
| REGULATION OF CELL CYCLE PROCESS | 0.001091137356 | 1.4727 | 185 |
| LOCOMOTION | 0.000000000488 | -1.6461 | 202 |
| DEFENSE RESPONSE TO SYMBIONT | 0.000000000000 | -1.8815 | 176 |
| INFLAMMATORY RESPONSE | 0.000000000000 | -1.9227 | 151 |
| CELL CHEMOTAXIS | 0.000000000721 | -2.2459 | 58 |
| HUMORAL IMMUNE RESPONSE | 0.000000000003 | -2.5619 | 47 |

**TableS5.** Significantly enriched Gene Ontology Biological Process (GOBP) pathways identified by gene set enrichment analysis under acute and chronic coal dust exposure conditions.

| **Antibody** | **Company** | **Cat. No.** | **Dilution** |
| --- | --- | --- | --- |
| Anti-ZO-1 Antibody | Rockland | 600-401-GU7 | 1:400 |
| VE-Cadherin | Cell Signaling Technology | 2500 | 1:200 |
| Cleaved Caspase-3 Antibody | Cell Signaling Technology | 9661 | 1:400 |
| Phospho-Histone H2A.X Antibody | Cell Signaling Technology | 2577 | 1:200 |
| Anti-Tubulin, Acetylated Antibody | Sigma-Aldrich | T6793 | 1:200 |

**Table S6.** Primary antibodies used for immunofluorescence staining.
